# Electrocorticographic Detection of Speech Networks in Glioma-infiltrated Cortex

**DOI:** 10.1101/2025.07.04.663249

**Authors:** Vardhaan S. Ambati, Amit Persad, Jasleen Kaur, Sanjeev Herr, Emily Cunningham, Abraham Dada, Justyna O. Ekert, Quinn Greicius, Alexander Silva, Paul M. Villalobos, Niels Olshausen, Youssef Sibih, Sena Oten, Hunter S. Yamada, Gray Umbach, Alexander Aabedi, Wajd Al-Holou, Jacob Young, Madhumita Sushil, Edward Chang, Mitchel S. Berger, David Brang, Shawn L. Hervey-Jumper

**Author notes:** **Corresponding Author**: Shawn L. Hervey-Jumper, MD, Department of Neurological Surgery, University of California, San Francisco, 513 Parnassus Ave, HSE-860, San Francisco, CA 94143-0112, David Brang, PhD, Department of Psychology, University of Michigan, 1004 East Hall, 530 Church St, Ann Arbor, MI 48109.

## Abstract

Direct cortical stimulation (DCS) is the clinical gold standard for identifying functional cortex in the human brain, which is essential for the safe removal of brain lesions. Defining the electro-physiological properties of DCS− positive cortical regions may facilitate the identification of critical language regions, thereby permitting safe glioma resections in communities without access. Leveraging a multicenter electrophysiologic dataset of DCS− positive language regions spatially matched with subdural arrays, we analyzed regions identified as functionally critical (DCS+) versus functionally non-critical (DCS–) during intraoperative language mapping. In IDH-mutant gliomas, DCS+ regions exhibited significantly greater speech-related neural activity and enhanced encoding and decoding of linguistic and semantic features. We demonstrate that resting-state classifiers distinguish DCS+ from DCS– regions in IDH-mutant tumors. Task-based and resting-state electrophysiologic distinctions were pathology-specific and not present in IDH-wildtype glioblastomas. These findings may accelerate DCS mapping by guiding surgeons to priority regions, improving efficiency, and patient outcomes.

## MAIN TEXT

Diffuse gliomas are the most prevalent primary brain tumors in both pediatric and adult populations, accounting for approximately 85% of cancer-related cognitive and behavioral deficits.^1,2^ A key factor contributing to these impairments is the disruption of neural networks underpinning language, sensorimotor functions, and cognition. The precise cortical locations and neural substrates involved in speech and cognitive processes vary substantially among individuals and are further altered by tumor infiltration.^3–9^ Achieving maximal safe resection of brain tumors like diffuse gliomas is the primary step in treatment, with nearly 500,000 such procedures performed globally each year.^10,11^ However, individual differences in cortical organization present substantial challenges to the safe excision of these tumors.^12,13^ Inadvertent damage to language-critical regions during surgery may cause persistent neurological deficits, adversely affecting quality of life and decreasing survival rates.^14–16^

For over 150 years, direct cortical stimulation (DCS) has been the gold standard method for identifying language-critical areas of the cortex. Initially developed by Fritsch and Hitzig in 1870 and further advanced by Wilder Penfield during the 1930s, this technique is primarily employed during awake craniotomies to map eloquent brain regions.^8,17–20^ Regions where electrical stimulation causes temporary speech disruptions are designated as DCS-positive (DCS+), and by tradition, these zones are spared during surgical resection to preserve language function. This approach has proven effective in maintaining neurological integrity and reducing postoperative deficits.^8,21^ Although the core technology behind DCS and brain mapping has remained essentially unchanged over the decades, it remains technically complex and resource-intensive, which limits its global implementation. Interestingly, consistent results obtained with various mapping techniques imply that the electrophysiological properties of the cortex are intimately related to the organization of language networks. However, the effects of DCS on language function, and the mechanisms by which these effects arise, are not well understood, and must be complemented with additional electrophysiological analysis of the targeted area of cortex before conclusions about its functional role can be made.^18^ Previous work on the electrophysiological properties of DCS+ regions has focused chiefly on epilepsy patients or non-tumor-infiltrated cortex in glioma cases.^22–24^ Recent ECoG studies indicate that gliomas do not simply cause tissue destruction but remodel neural circuits,^3,4,25,26^ suggesting that language-related DCS+ areas within infiltrated cortex may exhibit distinct electrophysiological characteristics.

In this study, we utilize a prospectively gathered, multi-center dataset consisting of subdural array local field potentials recorded during intraoperative awake ECoG brain mapping in patients. Our goal is to identify neural signatures that differentiate DCS+ from DCS– regions within glioma-infiltrated cortex. These insights could support the development of scalable tools that either complement DCS mapping or serve as independent, non-DCS-based approaches for identifying cortical language regions during surgery. We begin by analyzing the electrophysiological activity of DCS+ and DCS– regions during speech production. Next, we explore whether DCS+ areas serve as cortical information hubs. Lastly, we evaluate the performance of machine learning algorithms trained on resting-state, passive ECoG data in a training cohort to classify DCS+ regions in an external validation set accurately.

## RESULTS

### Language Mapping and Electrocorticography Cohort

Our prospective cohort included 81 consecutive patients with WHO 2–4 diffuse gliomas (per WHO 2021 criteria) who underwent awake language and sensorimotor mapping via direct cortical stimulation (DCS) with concurrent subdural electrocorticography (Fig. 1a). The mean age was 51 years (range 19–78), with 69.1% male; 77.8% had newly diagnosed gliomas. All patients underwent standard tumor resection in the language-dominant hemisphere, predominantly in the left hemisphere (95.1%) (Table 1, Fig. 1a, Extended Data Fig. 1a).

**Figure 1:**
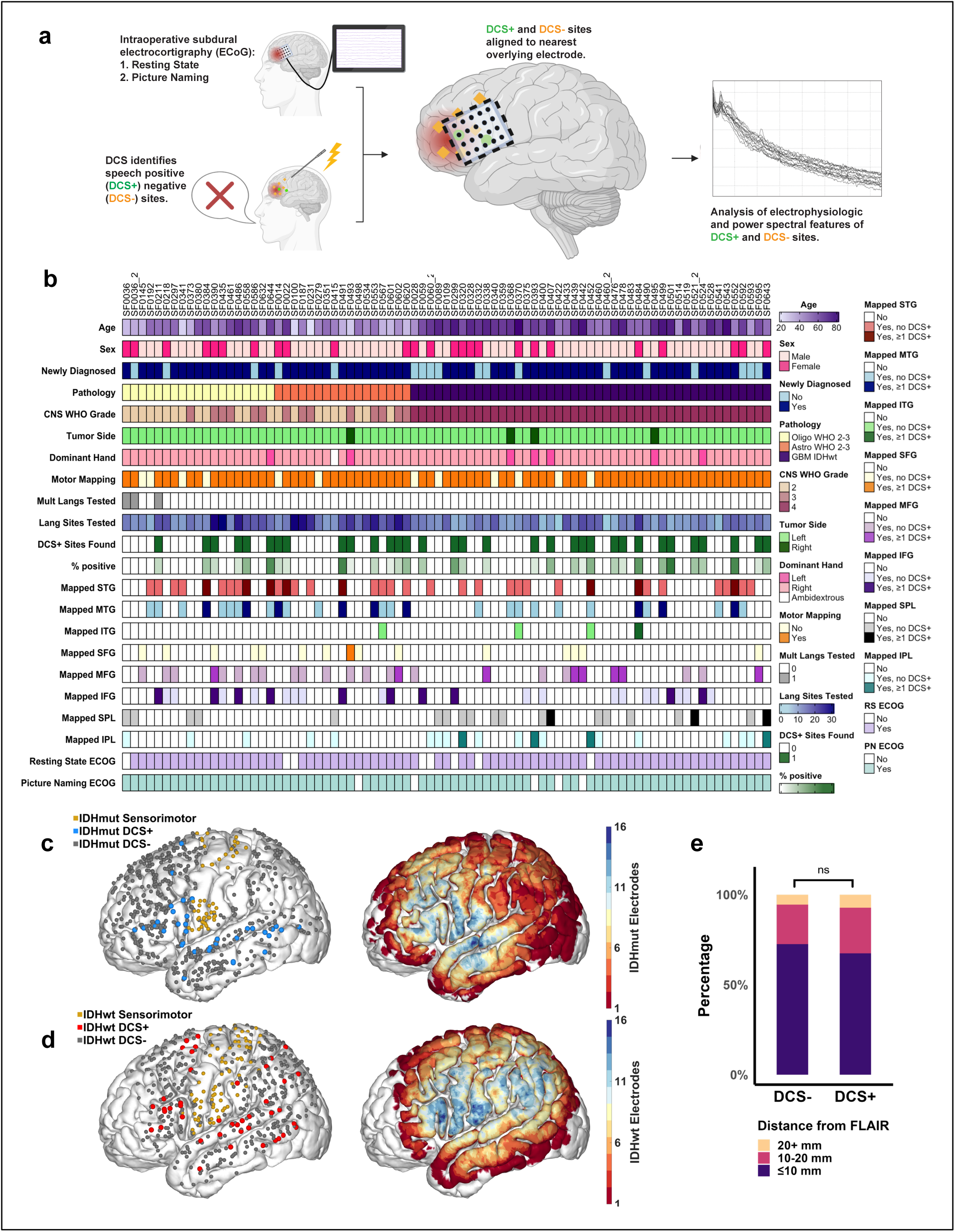
Electrode arrays and cortical regions showing positive language responses to direct stimulation. **a,** Schematic of the study design. Patients underwent intraoperative direct cortical stimulation (DCS) mapping to identify language sites within glioma-infiltrated cortex. High-density electrode arrays were placed to record human local field potentials from gliomas projecting to the cortical surface during the resting state and a semantic naming (picture naming) task. Mapped DCS positive (DCS+) sites were co-registered with their corresponding electrodes for subsequent electrophysiologic analysis. **b,** Oncoprint summarizing demographic, clinical, and mapping details for each participant in the study cohort (81 patients). **c-d,** Distribution of all mapped DCS language and motor sites in patients with (**c**) IDH-mutant (36 patients; 39 DCS+ sites; 576 DCS– sites) and (**d**) IDH-wildtype (45 patients; 44 DCS+ sites; 479 DCS– sites) gliomas, visualized on a standardized Montreal Neurological Institute (MNI) brain template. Heatmaps on MNI templates illustrate the spatial layout of electrode positions across patients in each cohort. **e,** DCS+ and DCS– sites do not differ significantly in distance from the FLAIR hyperintensity edge. Statistical significance was assessed using a Chi-squared test across distance bins (p = 0.58) and a two-sided linear mixed-effects model with patient-level random effects (p = 0.32). In panels **c** and **d**, DCS regions represent locations intended to be approximate rather than exact, as they were mapped on patient-specific pial surfaces and transformed into MNI space to coalesce data across patients. While true, precise DCS sites were used for analyses within each patient’s native space, the group maps in MNI space are approximate.

**Table 1.**

| **Variable** | **Study Cohort N=81** |
| --- | --- |
| Age, mean (range) | 51.0 (19.0, 78.0) |
| Sex, Male (%) | 56 (69.1%) |
| Newly Diagnosed (%) | 63 (77.8%) |
| Pathology |  |
| GBM IDHwt (%) | 45 (55.6%) |
| Oligodendroglioma WHO 2-3 (%) | 19 (23.5%) |
| Astrocytoma WHO 2-3 (%) | 17 (21.0%) |
| Dominant Hand |  |
| Right (%) | 73 (90.1%) |
| Left (%) | 7 (8.6%) |
| Ambidexterous (%) | 1 (1.2%) |
| Left Hemisphere Tumor (%) | 77 (95.1%) |
| Bilingual (%) | 10 (12.3%) |
| Motor Mapping Performed (%) | 69 (85.2%) |
| Multiple Languages Tested (%) | 3 (3.7%) |
| Number of Language Sites Tested, mean (range) | 14.0 (4.0, 31.0) |
| ≥1 DCS+ Found (%) | 33 (40.7%) |
| Number of Positive Sites per Patient, mean (range) | 1.0 (0.0, 7.0) |
| Percent Positive Sites per Patient, mean (range) | 7.0 (0.0, 42.9) |
| IFG, n = 27 |  |
| Mapped No DCS+ (%) | 17 (63.0%) |
| Mapped ≥1 DCS+ (%) | 10 (37.0%) |
| MFG, n = 26 |  |
| Mapped No DCS+ (%) | 18 (69.2%) |
| Mapped ≥1 DCS+ (%) | 8 (30.8%) |
| SFG, n = 15 |  |
| Mapped No DCS+ (%) | 14 (93.3%) |
| Mapped ≥1 DCS+ (%) | 1 (6.7%) |
| STG, n = 35 |  |
| Mapped No DCS+ (%) | 27 (77.1%) |
| Mapped ≥1 DCS+ (%) | 8 (22.9%) |
| MTG, n = 29 |  |
| Mapped No DCS+ (%) | 20 (69.0%) |
| Mapped ≥1 DCS+ (%) | 9 (31.0%) |
| ITG, n = 4 |  |
| Mapped No DCS+ (%) | 3 (75.0%) |
| Mapped ≥1 DCS+ (%) | 1 (25.0%) |
| IPL, n = 20 |  |
| Mapped No DCS+ (%) | 16 (80.0%) |
| Mapped ≥1 DCS+ (%) | 4 (20.0%) |
| SPL, n = 20 |  |
| Mapped No DCS+ (%) | 17 (85.0%) |
| Mapped $\geq 1$ DCS+ (%) | 3 (15.0%) |
| ECOG_Resting_State (%) | 73 (90.1%) |
| ECOG_Picture_Naming (%) | 76 (93.8%) |

Given the prognostic impact of IDH mutation status on cognition and survival, patients were categorized into IDH-mutant (n=36; 39 DCS+ regions, 576 DCS– regions) and IDH-wildtype (n=45; 44 DCS+ regions, 479 DCS– regions) cohorts (Fig. 1b-d, Table 1). Overall, 33 patients (40.7%) had at least one DCS+ language or motor site, with similar proportions in each cohort (IDH-mutant: 14/36; 38.9%; IDH-wildtype: 19/45; 42.2%). The wildtype group was significantly older (58.9[±[11.7 vs. 41.1[±[12.8 years, p<0.001) and had fewer language regions tested per patient (11.6[±[3.9 vs. 17.1[±[6.8, p<0.001). No significant differences were observed between cohorts concerning sex, primary versus recurrent disease, tumor laterality, or the percentage of motor and DCS+ language regions (Table 2).

**Table 2.**

| Variable | IDH-mutant<br>N=36 (44.4%) | IDH-wildtype<br>N=45 (55.6%) | p value |
| --- | --- | --- | --- |
| Age, mean (range) | 41.1 ± 12.8 | 58.9 ± 11.7 | <0.001 |
| Sex, Male (%) | 25 (69.4%) | 31 (68.9%) | 0.957 |
| Newly Diagnosed (%) | 31 (86.1%) | 32 (71.1%) | 0.107 |
| Pathology |  |  | <0.001 |
| GBM IDHwt (%) | 0 (0.0%) | 45 (100.0%) |  |
| Oligodendroglioma WHO 2-3 (%) | 19 (52.8%) | 0 (0.0%) |  |
| Astrocytoma WHO 2-3 (%) | 17 (47.2%) | 0 (0.0%) |  |
| Dominant Hand |  |  | 0.334‡ |
| Right (%) | 33 (91.7%) | 40 (88.9%) |  |
| Left (%) | 2 (5.6%) | 5 (11.1%) |  |
| Ambidexterous (%) | 1 (2.8%) | 0 (0.0%) |  |
| Left Hemisphere Tumor (%) | 35 (97.2%) | 42 (93.3%) | 0.625‡ |
| Bilingual (%) | 7 (19.4%) | 3 (6.7%) | 0.1‡ |
| Motor Mapping Performed (%) | 30 (83.3%) | 39 (86.7%) | 0.675 |
| Multiple Languages Tested (%) | 3 (8.3%) | 0 (0.0%) | 0.084‡ |
| Number of Language Sites Tested, mean (range) | 17.1 ± 6.8 | 11.6 ± 3.9 | <0.001 |
| ≥1 DCS+ Found (%) | 14 (38.9%) | 19 (42.2%) | 0.762 |
| Number of Positive Sites per Patient, mean (range) | 1.1 ± 1.7 | 1.0 ± 1.6 | 0.779 |
| Percent Positive Sites per Patient, mean (range) | 5.9 ± 9.8 | 7.8 ± 11.4 | 0.428 |
| IFG (%), n = 27 |  |  | 0.695‡ |
| Mapped No DCS+ (%) | 8/14 (57.1%) | 9/13 (69.2%) |  |
| Mapped ≥1 DCS+ (%) | 6/14 (42.9%) | 4/13 (30.8%) |  |
| MFG (%), n = 26 |  |  | 0.038‡ |
| Mapped No DCS+ (%) | 13/15 (86.7%) | 5/11 (45.5%) |  |
| Mapped ≥1 DCS+ (%) | 2/15 (13.3%) | 6/11 (54.5%) |  |
| SFG (%), n = 15 |  |  | 1.0‡ |
| Mapped No DCS+ (%) | 9/10 (90.0%) | 5/5 (100.0%) |  |
| Mapped ≥1 DCS+ (%) | 1/10 (10.0%) | 0/5 (0.0%) |  |
| STG (%), n = 35 |  |  | 0.7‡ |
| Mapped No DCS+ (%) | 14/19 (73.7%) | 13/16 (81.2%) |  |
| Mapped ≥1 DCS+ (%) | 5/19 (26.3%) | 3/16 (18.8%) |  |
| MTG (%), n = 29 |  |  | 0.694‡ |
| Mapped No DCS+ (%) | 11/17 (64.7%) | 9/12 (75.0%) |  |
| Mapped ≥1 DCS+ (%) | 6/17 (35.3%) | 3/12 (25.0%) |  |
| ITG (%), n = 4 |  |  | 1.0‡ |
| Mapped No DCS+ (%) | 1/1 (100.0%) | 2/3 (66.7%) |  |
| Mapped ≥1 DCS+ (%) | 0/1 (0.0%) | 1/3 (33.3%) |  |
| IPL (%), n = 20 |  |  | 0.53‡ |
| Mapped No DCS+ (%) | 5/5 (100.0%) | 11/15 (73.3%) |  |
| Mapped ≥1 DCS+ (%) | 0/5 (0.0%) | 4/15 (26.7%) |  |
| SPL (%), n = 20 |  |  | 0.539‡ |
| Mapped No DCS+ (%) | 5/5 (100.0%) | 12/15 (80.0%) |  |
| Mapped ≥1 DCS+ (%) | 0/5 (0.0%) | 3/15 (20.0%) |  |
| ECOG_Resting_State (%) | 33 (91.7%) | 40 (88.9%) | 0.727‡ |
| ECOG_Picture_Naming (%) | 35 (97.2%) | 41 (91.1%) | 0.375‡ |

‡Fisher Exact Test

The entire cohort included 4,824 ECoG electrodes across 81 patients. In the IDH-mutant group (n=2,228 electrodes), electrode locations are depicted as a group-level heatmap (Fig. 1c) and individual positions (Extended Data Fig. 1b). The IDH-wildtype cohort (n=2,596 electrodes) is similarly shown with a heatmap (Fig. 1d) and electrode locations (Extended Data Fig. 1c). All DCS+ regions were situated within or adjacent to MRI-visible FLAIR hyperintense regions, consistent with glioma infiltration. No significant spatial differences were observed between DCS+ and DCS– regions across mutant and wildtype cohorts (Fig. 1e; Extended Data Fig. 1d–e). Electrode and DCS cortical region alignments are detailed in Extended Data Fig. 1f-g. A breakdown of DCS region-to-electrode mappings for each cohort, along with their inclusion in the picture naming and resting state analysis, is shown in Extended Data Fig. 2a-b. The number of DCS+ and DCS– regions included in each task-specific analysis is based on electrode coverage. Task-specific retained electrodes are described in each corresponding section below.

### Differential, Pathology-Dependent Activation of DCS+ and DCS*–* Regions

Previous studies have demonstrated bidirectional interactions between neurons and glioma cells, with neuronal activity driving glioma growth and gliomas increasing neuronal excitability. Diffuse gliomas remodel neural circuitry such that task-relevant neural responses activate tumor-infiltrated cortex.^3,4^ We therefore set out to determine whether DCS+ and DCS– regions differ in local coordination and activation during semantic naming, specifically picture naming. In the IDH-mutant cohort (n = 35), after preprocessing and alignment, 31 DCS+ and 342 DCS– electrodes were analyzed (Fig 2a, Extended Data Fig. 2a-b). Six DCS+ and 40 DCS– electrodes in the superior temporal gyrus—due to their stimulus-locked dynamics—were assessed separately, while the remaining electrodes contributed to broadband and high-gamma analyses (Fig. 2b-e; Extended Data Fig. 3b).

**Figure 2.**
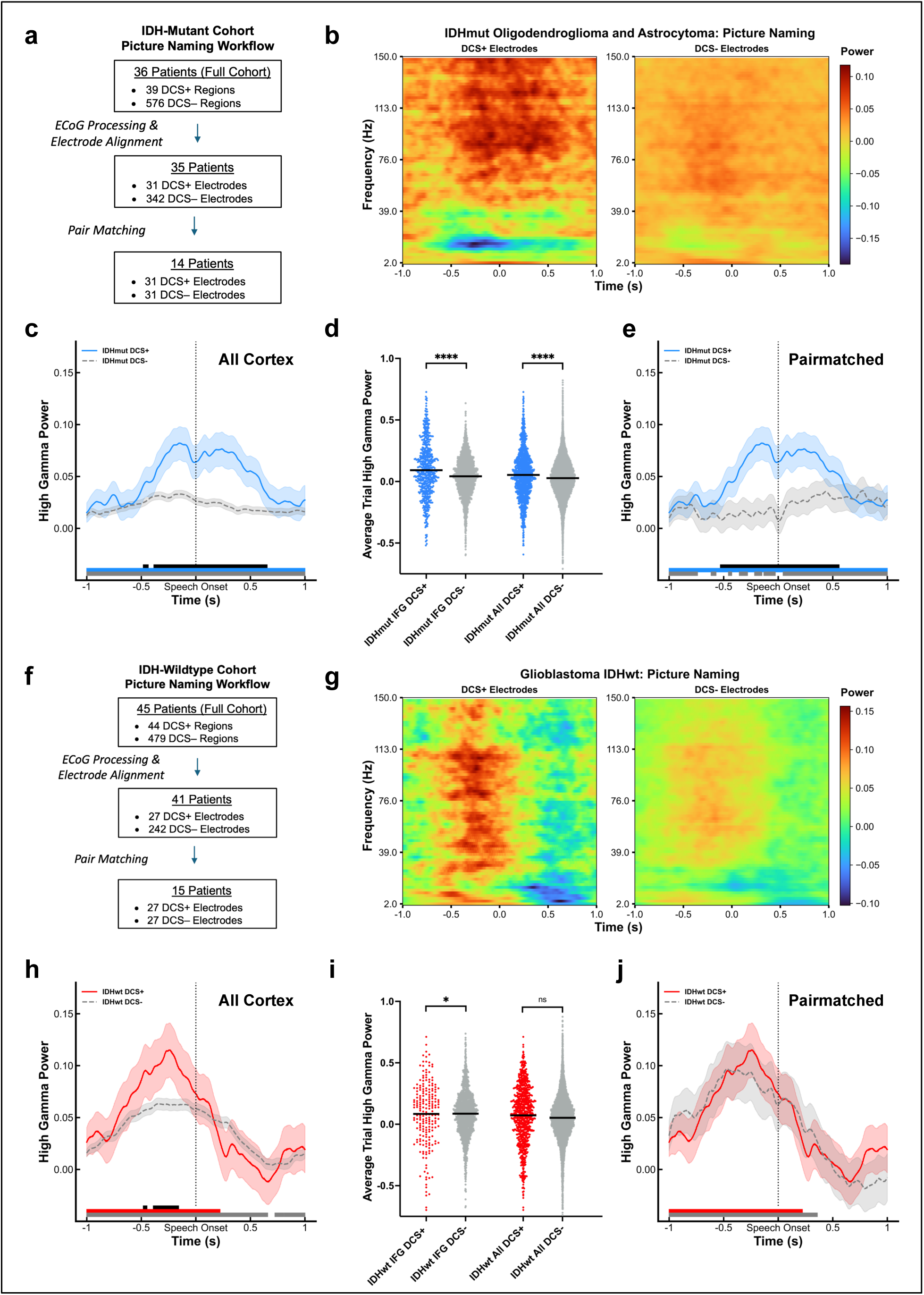
Direct cortical stimulation positive regions exhibit pathology-dependent event-related spectral perturbations. **a–e**, IDH-mutant oligodendroglioma and astrocytoma cohort with DCS+ and DCS– picture naming responses (35 patients). **a.** Flow chart for the IDHmut picture naming task showing the total number of patients and DCS sites, followed by the number of patients and sites after ECoG processing and electrode alignment, and finally after DCS+ to DCS– pair-matching and cleanup. **b**, Time–frequency spectra illustrate broadband (2–150 Hz) effects (differences from baseline [−1.2 to −1.0 seconds prior to speech onset] at DCS+ electrodes (25 electrodes; 1029 electrode-trial pairs) and DCS– electrodes (302 electrodes; 13120 electrode-trial pairs) during speech production. Speech onset was at time 0 seconds. Clear differences can be observed in the high-gamma range, most evident at frequencies above 70 Hz. **c**, High-gamma (70–120 Hz) event-related spectral perturbations (ERSPs) reveal significantly greater activation at DCS+ sites than DCS– sites. Timepoints with significant differences between DCS+ and DCS− sites (after FDR correction) are indicated by black bars. Colored bars represent time points where each cohort’s High-gamma activation was significantly higher than its baseline period (−1.2 to −1.0 s before speech onset, blue line represents DCS+ compared to baseline, and gray line represents DCS− compared to baseline). **d**, In the IDH-mutant cohort, the mean high-gamma power per trial (−0.5 seconds prior to speech onset to 0.5 seconds after onset) per electrode is grouped by DCS status. DCS+ sites show significantly higher High-gamma power in the inferior frontal gyrus (IFG DCS+ 0.092 ± 0.221 vs. IFG DCS– 0.042 ± 0.158, *p* < 0.001) and across all cortical regions (all DCS+ 0.054 ± 0.120 vs all DCS– 0.027 ± 0.158, *p* < 0.001). **e**, Pair-matched comparison (14 patients, 31 DCS+, 31 DCS–) shows sustained, significantly greater High-gamma activation at DCS+ sites, with significant time points indicated by bars at the bottom of the plot as in (**b**). **f–j**, IDH-wildtype glioblastoma cohort with DCS+ and DCS– picture naming responses (41 patients). **f**, Flow chart for the IDHwt picture naming task showing the total number of patients and DCS sites, followed by the number of patients and sites after ECoG processing and electrode alignment, and finally after DCS+ to DCS– pair-matching and cleanup. **g**, Time–frequency spectral data demonstrate broadband (2–150 Hz) at IDH-wildtype DCS+ (20 electrodes; 646 electrode-trial pairs) and DCS– (217 electrodes; 8027 electrode-trial pairs) sites during speech production. Speech onset was at time 0 seconds. **h**, High-gamma ERSPs show significantly greater activation at DCS+ (20 electrodes; 646 electrode-trial pairs) than DCS– electrodes (216 electrodes; 8027 electrode-trial pairs). **i**, Mean High-gamma power per trial per electrode reveals significantly lower high-gamma power at DCS+ sites in the IFG (IFG DCS+ 0.084 ± 0.260 vs. IFG DCS– 0.086 ± 0.188, *p* = 0.022); however, no differences are observed across all cortical regions (all DCS+ 0.073 ± 0.24 vs all DCS– 0.052 ± 0.188, *p* = 0.065). **j**, Pair-matched (13 patients, 27 DCS+, 27 DCS–) comparison reveals no significant high-gamma differences between IDH-wildtype DCS+ and DCS– sites. In panels **c**, **e**, **h**, and **j**, solid lines represent the cohort-averaged time series across trials and electrodes; shaded areas denote the standard error of the mean (SEM). Time series were grouped by DCS status and compared at each time point using Welch’s *t*-tests with false discovery rate (FDR) correction for multiple comparisons. Time points with significant differences are marked by black bars above the x-axis. Colored bars denote time points where each cohort’s High-gamma power was significantly greater than its baseline (paired *t*-test, FDR corrected). In panels **c** and **h**, horizontal bars indicate the group-level median of high-gamma power. Significance was assessed using two-tailed linear mixed-effects models controlling for patient-level random effects.

First, in the IDH-mutant cohort, we examined spectral activity over the broad range of 2-150 Hz during speech. Both DCS+ and DCS– electrodes showed clear deviations from baseline (−1.2 to −1.0 s pre-speech), particularly in the high gamma range (70-150 Hz) with DCS+ electrodes exhibiting more prominent, temporally structured responses (Fig. 2b). We focused on the high-gamma band (70–150 Hz), which reflects local neuronal firing and cortical hyperexcitability, and in this context, is aligned with speech onset.^27–29^ DCS+ electrodes demonstrated significantly greater and more sustained ERSPs than DCS– electrodes across the time series, as determined by Welch’s t-tests with false discovery rate (FDR) correction (Fig. 2c). Data were normalized to the average of the pre-stimulus baseline periods across all trials using the VSSUM approach: (Data - Baseline)/(Data + Baseline). Within the canonical speech area of the lateral prefrontal cortex (LPFC) in the inferior frontal gyrus (IFG),^30–32^ 14 DCS+ and 62 DCS– electrodes revealed task-related activity, with DCS+ electrodes showing enhanced responses (Extended Data Fig. 3a). Similar patterns were observed in the superior temporal gyrus (STG; 6 DCS+ and 40 DCS–), with signals aligned to stimulus onset, indicating consistent and generalizable behavioral response patterns within DCS+ regions (Extended Data Fig. 3b).

Within tumor-infiltrated cortex, task-evoked neural dynamics may reflect underlying pathological disruption, potentially distinguishing regions that retain functional relevance (DCS+) from those that do not (DCS–), as determined by direct cortical stimulation areas. We therefore sought to determine whether the magnitude and coordination of task-related neural activity within DCS+ regions differs from that of DCS− regions while accounting for regional variability and electrode distribution. Using linear mixed effects models to control for patient-level differences, we found that averaged high-gamma power per trial (−0.5 to +0.5 s relative to speech onset) was significantly higher at DCS+ electrodes compared to DCS– regions, notably in the IFG (DCS+: 0.092[±[0.221; DCS–: 0.042[±[0.158; p < 0.001). This pattern persisted across all regions (DCS+: 0.054[±[0.120; DCS–: 0.027[±[0.158; p < 0.001) (Fig. 2d). No statistically significant clinical or demographic differences existed between patients without pair-matched electrodes (all p > 0.05, Table 3, Extended Data Fig. 2c-d). To verify that these observations were not a product of regional variability/electrode distribution, we averaged high-gamma power across electrodes and pair matched each DCS+ electrode to its nearest DCS– electrode within the same gyrus and patient (14 patients; 31 pairs) (Fig 2a, Extended Data Fig. 2a-b). Pair-matched analysis confirmed elevated high-gamma responses at DCS+ electrodes (Fig. 2e).

**Table 3.**
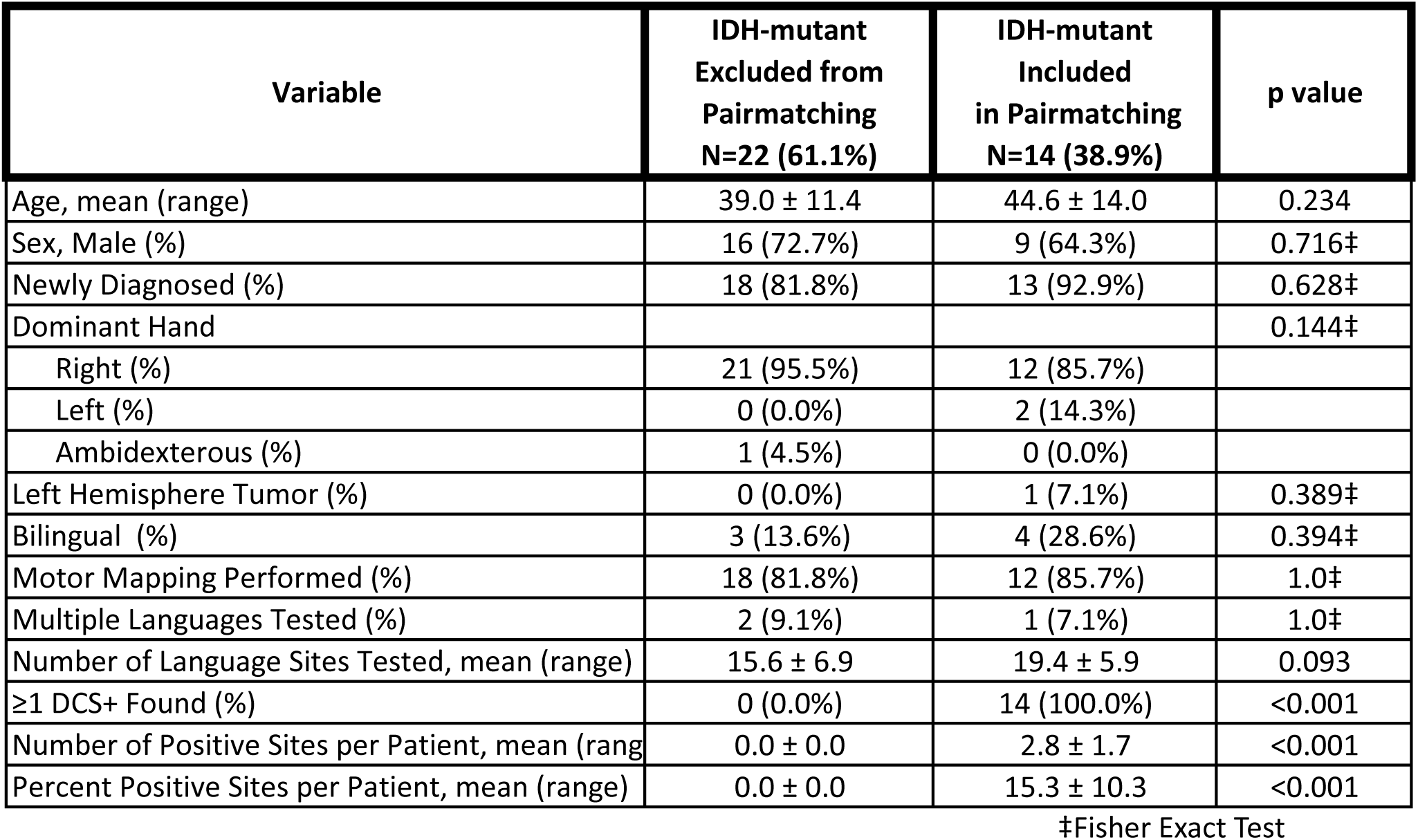

In the IDH-wildtype glioblastoma cohort (n=41), preprocessing yielded 27 DCS+ and 242 DCS– electrodes (Fig. 2f, Extended Data Fig. 2b). Seven DCS+ and 25 DCS– electrodes located in the STG were analyzed separately due to stimulus-locked dynamics (Extended Data Fig. 3b). Both groups showed robust broadband activity (2–150 Hz) during speech, with greater responses at DCS+ electrodes (Fig. 2g), and elevated high-gamma ERSPs (Fig. 2h). However, mean high-gamma power per trial reached statical significance in DCS+ electrodes in the IFG (0.084[±[0.260) compared to DCS– (0.086[±[0.188; p = 0.022), however with no significant difference when considering all regions (DCS+: 0.073[±[0.240; DCS–: 0.052[±[0.188; p = 0.065) (Fig. 2i). Furthermore, pair-matching within patients and gyrus (15 patients; 27 DCS+, 27 DCS–) revealed no significant high-gamma differences, indicating initial observations were likely regionally driven rather than intrinsic to DCS status (Fig. 2f, 2j, Extended Data Fig. 2a-b). No clinical or demographic differences were found between patients who were and were not included in the pair-matched analysis (all p > 0.05, Table 4).

**Table 4.**
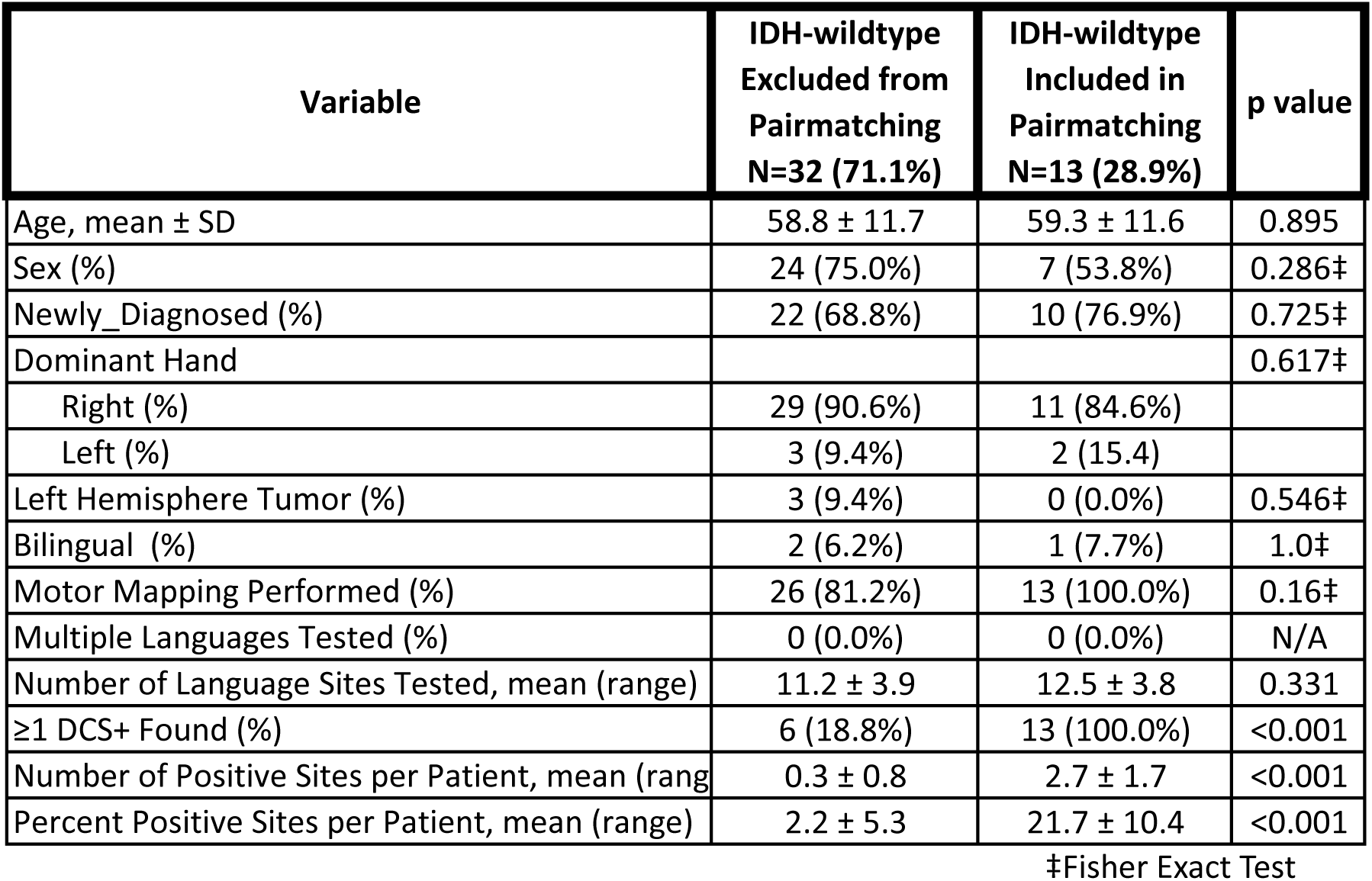

### Enhanced Information Representational Structure and Decoding at Behaviorally Critical Cortical Regions

In the clinical setting, DCS+ cortical regions are designated for preservation and are not resected, ensuring that this neural substrate remains with the patient throughout subsequent tumor-directed treatments, often for decades in patients with IDH-mutant tumors. It’s therefore possible that these cortically privileged regions contribute to cognitive processing, thereby providing a broader understanding of brain organization, information processing, and how neural circuits adapt in disease states. Having established differential task-specific broadband activity of DCS+ compared with DCS–IDH-mutant cortical regions, we sought to define the relative capacity of DCS+ and DCS− cortical regions to encode information. To ensure a rigorous within-subject comparison, analyses focused on pair-matched DCS+ and DCS– electrodes from the same patients. For each trial, we extracted high gamma power ERSPs from these electrodes and computed the Shannon entropy of the ERSP time series across trials at each time point.^4,33,34^ This provided a temporal profile of entropy values, serving as a statistical proxy for the diversity of neural states—and by extension, information encoding capacity—at each electrode.

In both the IDH-mutant and IDH-wildtype cohorts, DCS+ regions exhibited significantly higher entropy across the time series compared to their DCS– counterparts (IDH-mutant: *p* < 0.001; IDH-wildtype: *p* < 0.001; Fig. 3a–b), suggesting that DCS+ electrodes support a broader range of dynamic neural states. This effect remained when comparing DCS+ electrodes to all available DCS– electrodes (IDH-mutant: *p* < 0.001; IDH-wildtype: *p* < 0.001; Extended Data Fig. 4a–b), confirming the generalizability of the entropy finding across sampled regions.

**Figure 3.**
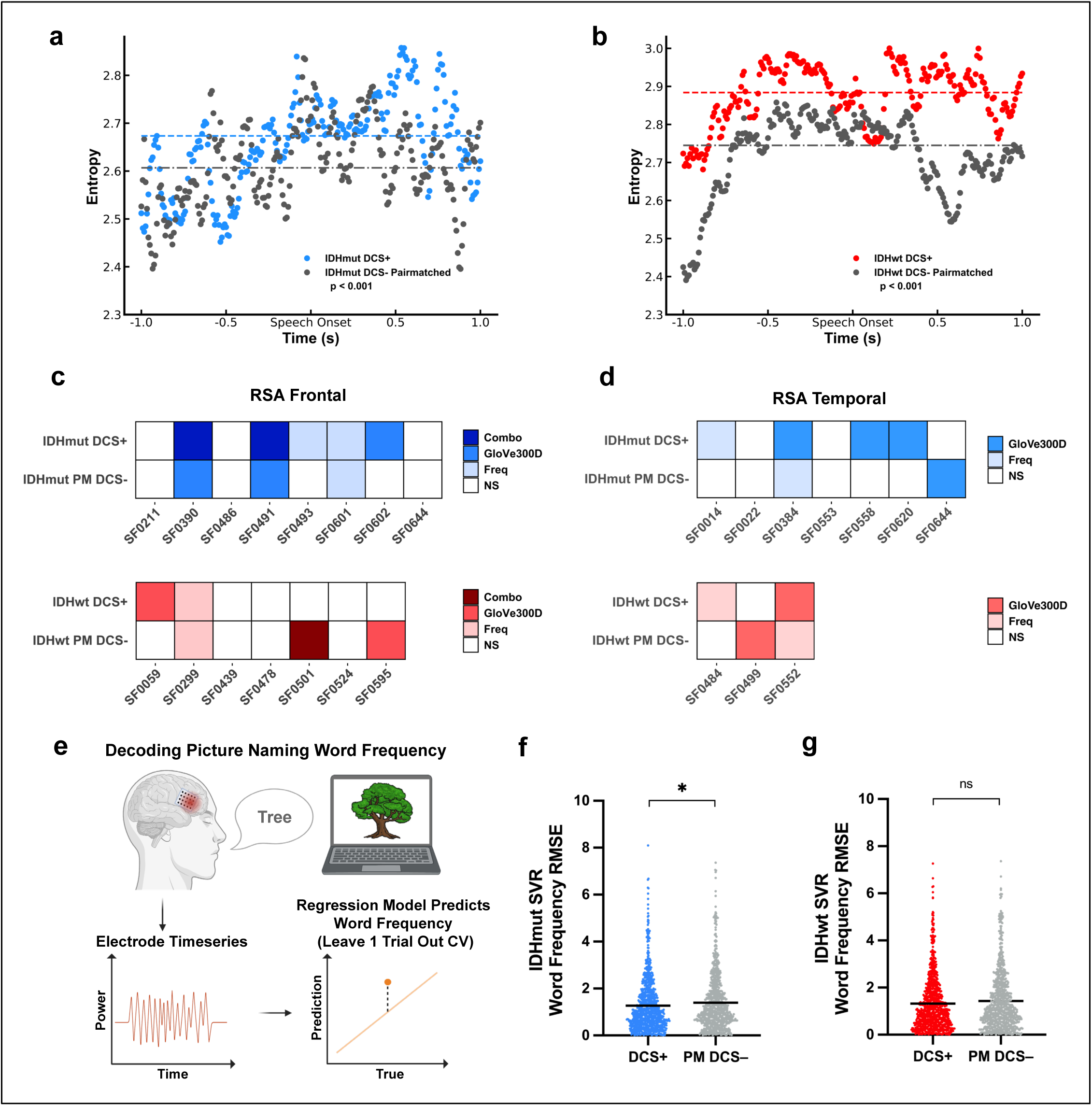
Information encoding and decoding accuracy of direct cortical stimulation positive regions. **a-b**, High gamma power event-related spectral potential (ERSP) entropy in **a**, IDH-mutant cohort, and **b**, IDH-wildtype cohort. ERSP timeseries from all trials were grouped by DCS status for each cohort and converted to group-level Shannon entropy of pair-matched electrodes at each time point. Entropy values were compared between DCS+ and DCS– regions using Welch’s t-tests. DCS+ sites exhibited significantly higher entropy in IDH-mutant (p < 0.001) and IDH-wildtype (p < 0.001) cohorts. Dashed lines indicate median entropy across all timepoints. Entropy values are displayed every five timepoints (approximately every 8.33 ms at 600 Hz). **c–d**, Heatmaps showing representational similarity analysis (RSA) across patients, with **c** representing the frontal and **d** the temporal lobes. Each column represents a patient, with the top row corresponding to each participant’s DCS+ electrodes and the bottom row to pair-matched DCS– electrodes. Shaded boxes indicate patients with at least one electrode showing significant RSA (blue). **c**, Frontal and **d,** Temporal lobes RSA was computed using word frequency, semantic, or both (combo) similarity from Global Vectors for Word Representation (GloVe) embeddings (300 dimensions). Across both regions, in the IDH-mutant cohort, DCS+ electrodes were more likely to show significant RSA within the same patient, suggesting increased encoding of linguistic features at DCS+ regions compared with DCS– regions. This trend was not apparent in the IDH-wildtype cohort (red). **e**, Schematic of the decoding task. Participants completed picture-naming trials, and multivariate regression models were trained to use high gamma time series data to predict the z-scored word frequency of the spoken word (derived from a large corpus using the wordfreq 3.1.1 Python package). Patient-level models were trained on all but one trial and tested on the held-out trial using cross-validation. Differences between predicted and true scores were used to compute trial-level root mean squared error (RMSE). RMSE values were then compared across electrodes grouped by DCS + or – status. **f**, In the IDH-mutant cohort, a support vector regression model trained on frontal lobe DCS+ electrodes (870 electrode-trial pairs), compared to pair-matched DCS– electrodes (776 electrode-trial pairs), demonstrated significantly lower trial-level RMSEs compared (DCS+: 1.27 ± 1.14 vs. DCS– : 1.40 ± 1.20, *p* = 0.030), indicating superior decoding performance. **g**, In the IDH-wildtype cohort, there was no significant difference in decoding RMSEs (DCS+: 1.38 ± 1.20 vs. DCS–: 1.48 ± 1.37, p=0.30) between frontal lobe DCS+ (368 electrode-trial pairs) and DCS– sites (368 electrode-trial pairs). **f–g**, Statistical significance was assessed using two-tailed linear mixed-effects models controlling for patient-level variability. Black bars indicate each cohort’s mean RMSE value.

Next, we assessed whether elevated entropy at DCS+ electrodes corresponded to structured neural representations using representational similarity analysis (RSA), comparing the trial-level cosine similarity of high-gamma time series with the linguistic features of the picture naming trial’s correct answer.^35^ In the frontal and temporal lobes, RSA incorporated z-scored word frequency and GloVe-based semantic similarity, reflecting prior evidence linking language embeddings of the picture naming trial’s correct answer to cortical representations.^36^ Analyses within individual patients compared DCS+ electrodes to pair-matched DCS– controls (Extended Data Fig. 4c). In the IDH-mutant cohort, DCS+ electrodes showed stronger RSA, compared to DCS– electrodes, with lexical and semantic features across both frontal and temporal regions, consistent across embedding dimensions (300- and 50-dimensional vectors), and replicated in extended analyses (Fig. 3c; Extended Data Fig. 4d). No such pattern was observed in the IDH-wildtype cohort (Fig. 3d; Extended Data Fig. 4d).

Finally, we asked whether increased entropy and representational structure at DCS+ regions translate into improved task decoding performance. For each electrode, we trained support vector regression models to predict word frequency from high-gamma time series and evaluated their performance using leave-one-trial-out cross-validation (Fig. 3e). In the IDH-mutant cohort, decoders trained on DCS+ electrodes yielded significantly lower root mean squared error (RMSE) compared to those trained on pair-matched DCS– electrodes (DCS+: 1.27[±[1.14 vs. DCS–: 1.40[±[1.20, *p* = 0.030; Fig. 3f), indicating superior decoding performance. Importantly, we confirmed that the decoding performance for both the DCS+ (DCS+ Shuffle: 2.60 ± 2.11, p<0.001) and DCS– (DCS– shuffle: 2.73 ± 2.22, p<0.001) electrodes was better than chance by shuffling the labels 10 times during each cross-validation fold (Fig. 3f, Extended Data Fig. 4e). However, in the IDH-wildtype cohort, no significant difference in decoding performance was observed between DCS+ and DCS– electrodes, using linear mixed effects models to control for patient-level differences (DCS+: 1.38[±[1.20 vs. DCS–: 1.48[±[1.37, p=0.30; Fig. 3g). While DCS+ and DCS– decoding performance in the IDHwt cohort did not differ, both DCS+ (DCS+ shuffle: 2.69 ± 2.38) and DCS– (DCS– shuffle: 2.48 ± 2.25) decoding was significantly better than their respective shuffles (p<0.001) (Extended Data Fig. 4f). These findings suggest that DCS+ regions in IDH-mutant glioma-infiltrated cortex support high-capacity, behaviorally relevant neural processing. They exhibit elevated entropy, enhanced representational structure, and improved linguistic decodability. These results may inform the development of future neuroprosthetics and brain-computer interfaces designed to decode speech from neural activity within glioma-infiltrated cortex.^37–40^ As neuroprosthetics are designed to interpret and decode behavior from neuronal activity, DCS+ regions may represent high-information-capacity nodes that could enhance decoding efficiency in IDH-mutant glioma patients.

### Resting State Prediction of DCS+ Regions in IDH-mutant Glioma

Awake brain mapping requires extensive resources, expertise, and infrastructure, limiting its availability at many hospitals worldwide. Recent studies have utilized resting-state connectivity features, such as connection strength, spectral power, and network metrics, to classify or predict DCS+ regions in nonmalignant epilepsy cortex.^41,42^ Diffuse gliomas remodel cortical speech representations; therefore, it remains unclear whether resting state spectral features can classify behaviorally relevant brain regions. Given the distinct electrophysiological differences between DCS+ and DCS– regions during speech tasks in IDH-mutant gliomas (Fig. 2), we examined whether intrinsic resting-state differences also exist. Analysis of PSD curves in 33 IDH-mutant patients (DCS+: 29 electrodes; DCS–: 278 electrodes) revealed clear separation between DCS+ and DCS– regions (40 patients; DCS+: 23; DCS–: 248) (Extended Data Fig. 2a-b, Fig 4a). This was absent in the IDH-wildtype cohort (Extended Data Fig. 5a). Restricting analysis to patients with both DCS+ and DCS– electrodes confirmed persistent differences in IDH-mutant (12 patients) but not in wildtype (12 patients) cases (Extended Data Fig. 5c–e).

**Figure 4:**
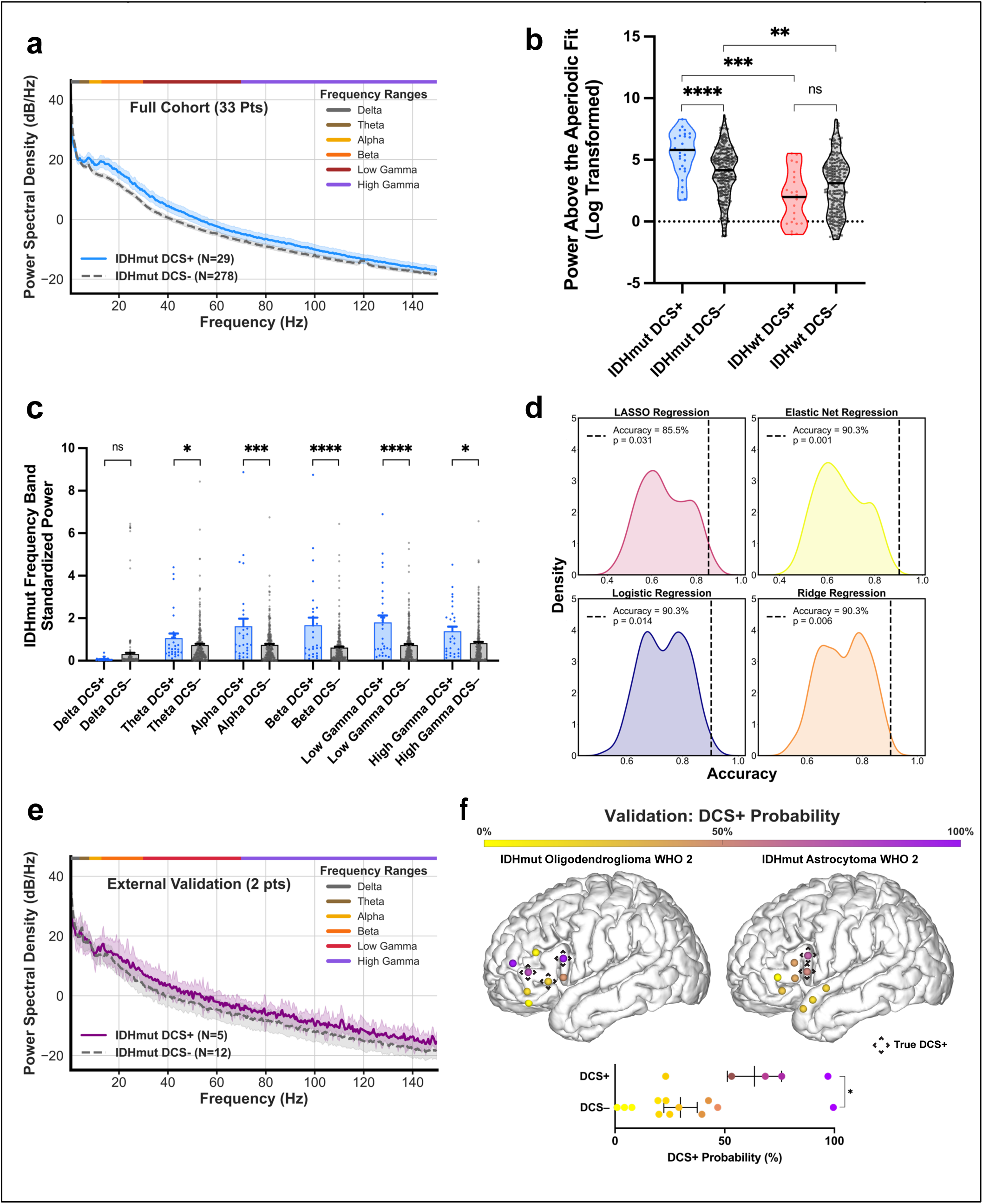
Resting state features predict direct cortical stimulation positive language regions in IDH mutant glioma. **a**, Power spectral density (PSD) curves from resting state recordings demonstrate separation of IDH-mutant electrodes (33 patients), including DCS+ (N=29 electrodes), and DCS– language regions (N=278 electrodes). Shaded regions indicate standard error of the mean (SEM). **b**, Mean power above aperiodic fit (aperiodic fit subtracted from PSD; arbitrary units) are shown for DCS+ and DCS– regions, stratified by IDH mutation status. In IDH-mutant tumors, DCS+ language sites exhibited significantly higher broadband power (5.44 ± 1.79) compared to DCS– sites (3.97 ± 2.07) (*p* < 0.001), while no such difference was observed in the IDH-wildtype group (IDHwt DCS+ 1.90 ± 2.12, IDHwt DCS– 2.89 ± 2.31) (*p* = 0.62). Across DCS+ sites, IDH-mutant cases had significantly higher broadband power than IDH-wildtype cases (*p* < 0.001); a similar pattern was observed at DCS– sites (*p* = 0.003). **c**, Standardized power in the Delta, Theta, Alpha, Beta, Gamma, and High Gamma frequency bands was compared between DCS+ and DCS– language regions in IDH-mutant gliomas. DCS+ sites exhibited significantly higher power across Theta (DCS+ 1.06 ± 1.19, DCS– 0.741 ± 0.978, *p* = 0.040), Alpha (DCS+1.62 ± 1.94, DCS– 0.749 ± 0.810, *p* < 0.001), Beta (DCS+ 1.67 ± 1.94, DCS– 0.618 ± 0.790, *p* < 0.001), Low Gamma (DCS+ 1.80 ± 1.75, DCS– 0.736 ± 0.830, *p* < 0.001), and High Gamma (DCS– 0.831 ± 0.968, DCS+ 1.39 ± 1.19, *p* = 0.011) frequency bands, with no significant differences observed in the Delta band (DCS+ 0.0602 ± 0.0777, DCS– 0.309 ± 1.05, *p* = 0.965). Bars represent the mean, and error bars represent the standard deviation. **d**, Results of non-parametric tests (1000 shuffles) for separate multivariate models, including Least Absolute Shrinkage and Selection Operator (LASSO), Elastic Net, Logistic, and Logistic with L2 regularization (Ridge) regression models. All models demonstrated high performance. LASSO accuracy- 0.855, AUCROC- 0.865, sensitivity- 0.571, specificity- 0.890; Elastic Net accuracy- 0.903, AUCROC- 0.813, sensitivity- 0.571, specificity- 0.946. Logistic Regression accuracy- 0.903, AUCROC- 0.746, sensitivity- 0.571, specificity-0.946. Ridge accuracy- 0.903, AUCROC- 0.746, sensitivity- 0.571, specificity- 0.946. Kernel density estimates (KDE) represent the distribution of shuffled accuracies, with a line indicating each model’s true accuracy and p-values indicating statistical significance. **e**, PSD from an external validation cohort (2 IDH-mutant patient; 5 DCS+, 12 DCS–) showing separation between DCS+ and DCS– electrodes, consistent with findings in Fig. 3a. **f**, Highlights validation cohort of two patients, 5 DCS+ and 17 DCS– regions, displayed on two separate MNI brains: the left brain represents an IDH-mutant oligodendroglioma (WHO grade 2) from the University of Michigan, and the right an IDH-mutant astrocytoma (WHO grade 2) from UCSF (different surgeon). Sites are colored according to a gradient based on the SoftMax probability scores of the best-performing regression model (LASSO), which achieved an accuracy of 88.2%, an AUROC of 0.817, a sensitivity of 0.80, and a specificity of 0.917. The model was trained on the entire study cohort (only including patients with ≥1 DCS+ sites) and tested on the validation cohort. DCS+ sites, indicated by a black dashed diamond outline, exhibit higher probability scores (more purple). A dot plot below, with dots also colored according to the SoftMax color gradient, demonstrates that DCS+ sites have significantly higher probability predictions (p = 0.032), as determined by a linear mixed effects model, controlling for patient-specific differences. **b,** The black line in each violin plot represents the group’s median value. **b-c**, Statistical significance was assessed using two-tailed linear mixed-effects models controlling for patient-level variability. **f**, Horizontal black lines on each dot plot represent the group’s mean probability and standard error of the mean

For these analyses, we quantified spectral power. We examined both overall activity beyond the aperiodic fit, as well as band-delimited activity within conventionally defined Delta (0.2–3.99 Hz), Theta (4–7.99 Hz), Alpha (8–12.99 Hz), Beta (13–29.99 Hz), Gamma (30–69.99 Hz), High Gamma (70–150 Hz) bands. In IDH-mutant tumors, DCS+ electrodes showed significantly higher broadband deviation than DCS– electrodes (5.44[±[1.79 vs. 3.97[±[2.07, p < 0.001; Fig. 4b) from the aperiodic fit, whereas no difference was observed in IDH-wildtype regions (1.90[±[2.12 vs. 2.89[±[2.31, p = 0.62) (Fig. 4b).^43,44^ IDH-mutant cortices exhibited greater broadband activity overall (p < 0.001) and showed significant increases in theta, alpha, beta, low gamma, and high gamma power at DCS+ electrodes (all p < 0.05), which suggests synchronized, physiologically structured signals (Fig. 4c).^43^ These differences were absent in IDH-wildtype electrodes. Concurrently, the higher high gamma power—commonly associated with increased local neuronal spiking— even at rest, may reflect elevated baseline cortical organization and coordination.

These results suggest that at rest, DCS+ regions in IDH-mutant gliomas retain structured oscillatory patterns and stable network dynamics relative to DCS– regions. These resting-state features that may aid in the cortex’s capacity to support or decode task-related activity, particularly for functions like speech and cognition. In contrast, DCS+/DCS– regions in IDH–wild-type glioblastoma do not differ significantly in resting state cortical disorganization, potentially impairing cognition-related functional decodability.

Given the power spectral density (PSD) and broadband activity differences associated with DCS status in IDH-mutant gliomas, we trained four multivariate classifiers—LASSO, Elastic Net, Ridge, and logistic regression—to predict DCS status from broadband PSD spectra in the IDH-mutant cohort. We trained and tested (testing cohort: 7 DCS+ electrodes; 55 DCS– electrodes) four multivariate linear models—LASSO, Elastic Net, Ridge regression, and logistic regression—within our study cohort. All models achieved high performance accuracies, ranging from 85.5% to 90.3%, AUROC values from 0.746 to 0.865, and high specificity (0.890–0.946); however, sensitivity was more modest, possibly due to data imbalance, with a value of 0.571 across all models. Specifically, LASSO achieved an accuracy of 0.855 (AUROC = 0.865, sensitivity = 0.571, specificity = 0.890); Elastic Net, 0.903 (AUROC = 0.813, sensitivity = 0.571, specificity = 0.946); logistic regression, 0.903 (AUROC = 0.746, sensitivity = 0.571, specificity = 0.946); and Ridge regression, 0.903 (AUROC = 0.746, sensitivity = 0.571, specificity = 0.946). All models significantly outperformed chance, as assessed by a non-parametric permutation test with 10,000 label shuffles (p < 0.001; Fig. 4d).

To evaluate model generalizability, we retrained each model on the full study cohort, limited to patients with both DCS+ and DCS– regions (to reduce data imbalance), and tested on an external validation consisting of 2 patients (1 IDH-mutant Oligodendroglioma WHO grade 2 [University of Michigan] and 1 IDH-mutant Astrocytoma WHO grade 2 [UCSF different surgeon]) together with 5 DCS+ and 12 DCS– electrodes. Importantly, the PSD of the IDH-mutant validation cohort replicated the spectral separation seen in Fig. 4a (Fig. 4e). The best-performing model, LASSO regression, achieved an accuracy of 88.2%, an AUROC of 0.817, a sensitivity of 0.80, and a specificity of 0.917. Importantly, SoftMax probability scores from this model were significantly higher for DCS+ regions compared to DCS– regions, as demonstrated by linear mixed-effects modeling controlling for patient differences (p = 0.032; Fig. 4f), indicating robust generalization and suggesting electrophysiologic models can reliably differentiate DCS status in IDH-mutant gliomas. The remaining models (Ridge, elastic net, and logistic regression) all also reported high accuracies (0.706-0.765), AUROCs (0.80-0.867), sensitivity (0.80 in all models), specificity (0.667-0.75), and significantly higher SoftMax scores for DCS+ regions (p<0.05) (Table 5). These findings suggest that resting-state classifiers may serve as a potential priority map to guide intraoperative stimulation, highlighting high-probability DCS+ regions to enable more efficient and targeted mapping.

**Table 5.**

| Model | Accuracy | AUROC | Sensitivity | Specificity | SoftMax p value |
| --- | --- | --- | --- | --- | --- |
| LASSO Regression | 0.882 | 0.817 | 0.8 | 0.917 | 0.032 |
| Logistic Regression | 0.765 | 0.867 | 0.8 | 0.75 | 0.005 |
| Elastic Net Regression | 0.706 | 0.8 | 0.8 | 0.667 | 0.045 |
| Ridge Regression | 0.765 | 0.867 | 0.8 | 0.75 | 0.005 |

## DISCUSSION

In 1870, Eduard Hitzig and Gustav Fritsch demonstrated that electrical stimulation of the canine motor cortex elicited limb movements, an essential discovery later extended to a human patient.^20^ These foundational findings established the principles underlying direct cortical stimulation (DCS) brain mapping via awake craniotomy procedures. Since its initial description in the 1930s, the DCS technique has maintained a consistent conceptual framework.^17^ Over subsequent decades, various stimulation parameters have been developed; however, despite methodological heterogeneity, DCS has proven to be a reliable method for localizing language-eloquent cortex across diverse clinical settings and patient populations.^45^ While the technological approach remains essentially unchanged, the neurophysiological underpinnings explaining why cortical regions within the dominant perisylvian cortex are critical for speech production are not yet fully elucidated.^8,18,32,46^ Despite its widespread clinical utility, DCS faces notable limitations, including risks of stimulation-induced seizures,^47^ the necessity for patient wakefulness during speech mapping, and the requirement for highly specialized multidisciplinary teams,^8^ which collectively render the procedure resource-intensive.

We conducted a multicenter study examining glioma-infiltrated cortex, incorporating electrophysiological assessments of regions annotated as DCS+ versus DCS– during language tasks. Our analysis revealed that electrophysiological distinctions are most prominent in IDH-mutant gliomas.^1^ Within this subgroup, DCS+ regions exhibited significantly elevated high-gamma (70-150 Hz) activity during speech initiation, indicative of increased local cortical activation. Additionally, these regions demonstrated higher Shannon entropy, reflecting greater information processing capacity, and superior encoding and decoding accuracy of linguistic and semantic speech features.^4^ These results suggest that, compared to DCS–regions, DCS+ regions in IDH-mutant gliomas may maintain or reorganize functional language networks, actively participating in language processing despite tumor infiltration of eloquent cortex.

Various neuroimaging and electrophysiological techniques have been proposed to complement or overcome the limitations of awake cortical mapping. Adjunctive modalities such as functional MRI (fMRI), transcranial magnetic stimulation (TMS), facial electromyography (EMG), and corticocortical evoked potentials (CCEPs) have been extensively studied.^32,48–51^ Nonetheless, none have superseded DCS in clinical practice, primarily due to their insufficient spatial accuracy in delineating cortical speech regions compared to the gold standard of intraoperative DCS mapping. Prior electrophysiological investigations in epilepsy and non-tumor-infiltrated cortex within glioma patients have revealed distinct spectral profiles and connectivity patterns.^22,24^ However, it remains unclear how tumor infiltration modulates these electrophysiological relationships—a crucial issue, since the primary goal of DCS in glioma surgery is to identify and preserve functionally critical infiltrated cortex. Our previous research indicates that gliomas remodel neuronal circuits, and that glioma-infiltrated cortex can exhibit synchronized activity, albeit with altered structural properties.^3,4^ These insights suggest that electrophysiologic differences between DCS+ and DCS– regions, potentially detectable via non-stimulation-based measures, may reflect underlying functional distinctions. This raises the possibility of passive, less invasive functional mapping approaches that rely on intrinsic electrophysiological signatures.

Physiological specialization of DCS+ regions, even within glioma-infiltrated cortex, carries significantly broader implications. Although IDH-mutant gliomas are less prevalent than IDH-wildtype tumors, their association with improved overall survival results in a higher long-term patient population.^1,11^ Consequently, a subset of these patients may experience persistent language deficits, highlighting the potential importance of physiologically characterized DCS+ regions as high-information cortical hubs. Such regions could be leveraged as targets for future brain-computer interface (BCI) applications designed to restore, compensate for, or augment language functions in individuals with tumor-related or post-treatment aphasia.^39,40,52^

In contrast, the lack of distinguishable electrophysiological features between DCS+ and DCS– regions in IDH-wildtype tumors raises critical biological and clinical questions. The findings of this study imply that even DCS+ regions within IDH-wildtype gliomas may lack the structured electrophysiologic signatures essential for robust language encoding. This may reflect disrupted language pathways or an environment characterized by glioblastoma-associated hyperexcitability that impairs typical cortical organization, though the precise mechanisms remain uncertain. Notably, recent investigations have demonstrated no significant difference in short-term neurological outcomes between awake and asleep glioblastoma resections.^53,54^ This contrasts sharply with data from low-grade glioma surgeries, where awake mapping has consistently shown benefits in functional preservation and increased extent of resection.^55,56^

This study is the first to demonstrate that resting-state electrocorticography (ECoG) can reliably distinguish between functionally eloquent and non-eloquent cortical regions in glioma patients. Specifically, we show that DCS+ regions in patients with IDH-mutant tumors exhibit increased broadband activity and frequency-specific oscillations—neural signatures previously linked to coordinated neuronal spiking and indicative of a healthier, more functionally intact cortex.^57–59^ Conversely, the absence of these electrophysiological signatures in glioblastoma (GBM) patients aligns with prior literature suggesting that high-grade gliomas are more aggressive and cause widespread cortical disorganization. In contrast, low-grade gliomas tend to infiltrate and integrate with surrounding tissue in a manner that preserves or permits remodeling of functional capacity, enabling DCS+ regions to maintain or recover their functional integrity.^3,60–65^ Machine learning models trained on power spectral features derived from resting-state recordings accurately classified DCS+ versus DCS– regions within our cohort and in an independent external dataset, despite differences in stimulation protocols. Notably, these models exhibited higher SoftMax probabilities for DCS+ regions in IDH-mutant patients, suggesting a graded, rather than binary, distinction, which underscores the potential clinical utility of probabilistic classification approaches.

These findings have significant translational implications. Resting-state electrophysiological classification could serve as a feasible adjunct or alternative to intraoperative awake speech mapping, particularly in patients with IDH-mutant gliomas. Prior studies have demonstrated that glioma-associated electrophysiologic signals are largely preserved across awake and anesthetized states, supporting the potential utility of resting-state mapping under anesthesia.^50^

Here we present NeuroGrid, an open-source, artificial intelligence (AI)-based diagnostic system for detecting language regions in IDH-mutant glioma-infiltrated cortex using subdrual ECoG (an interactive demo is available at https://ephys-dcs.replit.app/). Our results provide a strong rationale for future research into passive functional mapping conducted under general or light anesthesia, which could mitigate the cognitive load, reduce procedure duration, and diminish the risks associated with prolonged awake mapping. Additionally, evidence indicates that intraoperative real-time ECoG is feasible and may assist in maximizing tumor resection.^66,67^ This approach may also offer a valuable alternative for patients who are unable to undergo awake mapping due to cognitive, psychological, or anesthetic contraindications. Further, even in patients who are able to tolerate awake mapping, resting-state ECoG classifiers may aid targeted stimulation mapping, reducing strain to patients and improving surgical efficiency.

This study is the first to comprehensively demonstrate that electrophysiological features, measured during both task-based and resting states, uncover meaningful pathology-dependent differences between DCS+ and DCS– regions within glioma-infiltrated cortex. Our findings underscore the impact of tumor pathology on neural substrates of language encoding, establishing a foundation for the development of next-generation, minimally invasive methods for functional language mapping. Ultimately, these insights may enhance intraoperative language preservation strategies and facilitate the advancement of brain-computer interface (BCI) applications in glioma surgical management.

### Limitations

While our study has numerous strengths, including being one of the largest electrophysiologic analyses of glioma and DCS regions to date, certain limitations are inherent. We use entropy as a surrogate for information encoding capacity and representational similarity analysis (RSA) to approximate the encoding of lexical information; these are computational techniques that have been previously validated but are inherently dependent on methodological parameters. Most notably, due to the ethical implications of resecting functional language regions, our study does not include tissue-level or molecular data and is therefore limited to the analysis of electrophysiologic signals collected during surgery. Further, our external validation cohort consisted of two patient. However, this is consistent with prior studies in this electrophysiologic domain, which often include only single-digit patient numbers—underscoring that even limited external validation remains meaningful in this context,^22,66^ especially given our large internal cohort and that the validation cohort was conducted under different DCS paradigms.

## CONCLUSION

We demonstrate clear pathology-dependent differences in language encoding between IDH-mutant and IDH-wildtype tumors. Specifically, DCS+ regions in IDH-mutant gliomas exhibit enhanced high-gamma activity during speech. Furthermore, IDH-mutant DCS+ regions may serve as information-processing nodes, as they demonstrate superior encoding and decoding of linguistic features compared to DCS– regions. Importantly, in IDH-mutant gliomas, broadband power spectral features from resting-state ECoG robustly distinguish DCS+ from DCS– regions, highlighting the potential for passive, less invasive functional mapping. These findings underscore an IDH-dependent physiological specialization that may inform intraoperative language preservation and pave the way for novel brain-computer interface applications in neuro-oncology.

## METHODS

### Cohort Description and Inclusion Criteria

The study protocol was approved by the University of California, San Francisco Institutional Review Board (CHR-17-23215). All participants were informed of the study’s purpose and procedures and provided written consent prior to participation.

We investigated the electrophysiologic correlates of language regions identified as either positive or negative by direct cortical stimulation (DCS; stimulation protocol and classification criteria detailed below), using subdural electrocorticography (ECoG; detailed below). Eligible participants were adults (≥18 years of age) undergoing resection of newly diagnosed or recurrent, cortically projecting diffuse gliomas located in the dominant language hemisphere, as determined by preoperative magnetoencephalography (MEG).^68^ All patients underwent standard-of-care intraoperative awake language mapping with concurrent ECoG, recorded during resting state and/or visual confrontation naming (picture naming) task. Patients were excluded if they were younger than 18 years, had non-diffuse glioma pathology (ex: brain metastases), lacked intraoperative awake language mapping, or had no usable intraoperative ECoG recordings.

Tumors were classified according to the 2021 World Health Organization criteria and included IDH-mutant oligodendrogliomas (Grades 2–3), IDH-mutant astrocytomas (Grades 2–3), and IDH-wildtype glioblastomas.^1^ There were no specific inclusion or exclusion criteria based on the lobe or gyral location of the tumor; patients were included if the clinical team determined that resection with intraoperative language mapping was indicated and all other eligibility criteria were met. Grade 4 IDH-mutant astrocytomas were excluded due to their relative rarity.^1^ Given the known relevance of IDH mutation status to glioma pathophysiology, we grouped IDH-mutant oligodendrogliomas and astrocytomas into a single IDH-mutant group. We considered IDH-wildtype glioblastomas as a separate group.

In total, 81 consecutive patients met inclusion criteria, including 33 with IDH-mutant gliomas. All patients underwent awake intraoperative language mapping with concurrent ECoG. Demographic and clinical characteristics are summarized in Table 1 and Figure 1b.

### Surgical Technique and Anesthesia Wash Out

Participants were positioned and underwent craniotomy and durotomy for cortical exposure solely based on clinical indication and surgical team decision-making, as described in our prior study.^8^ In brief, during the initial surgical phase, prior to full cortical exposure durotomy, most patients (with some variation depending on clinical factors) were anesthetized using our previously published protocol consisting of midazolam (2 mg), fentanyl (50-100 μg), propofol (50-100 μg/kg/min), and remifentanil (0.05-0.2 μg/kg/min).^8^ Prior to durotomy, lidocaine was locally administered to the scalp and dura. Upon completion of durotomy and subdural exposure, all systemic anesthetics and sedatives were discontinued.^8^ Consistent with our earlier studies, participants then underwent a minimum 15 to 20-minute anesthesia washout period followed by standardized wakefulness testing prior to brain mapping to ensure necessary arousal for reliable intraoperative language mapping and electrophysiologic recordings.^3,4^

### Intraoperative Language Mapping with Direct Cortical Stimulation

Discovery Cohort (UCSF): Before surgery, patients underwent comprehensive language assessments and were familiarized with the picture-naming task 1–2 days prior to the procedure. Only prompts that the patient answered correctly were utilized during language mapping. Direct cortical stimulation DCS was performed using a bipolar electrode stimulator with 5-mm spacing between contacts. Stimulation consisted of 1.25-millisecond biphasic square-wave pulses delivered in 4-second trains at 50-60 Hz, with a typical current intensity of 3–4 mA. On average, 14 language regions were tested per patient, each spaced approximately 1 cm apart. During each picture-naming trial, stimulation was applied to the cortical regions while the patient attempted to answer the picture-naming trial. Each region was tested with either 3 or 5 trials, and was classified as DCS-positive if stimulation resulted in errors in at least 2 of 3 or 3 of 5 trials, respectively. Errors included anomia, speech arrest, incorrect responses, or significant delays. Although not the primary focus of this study, many patients also underwent motor and sensory mapping using the same stimulation parameters.

Validation Cohort: Patients in the validation cohort underwent a direct cortical stimulation (DCS) paradigm that differed slightly from the training cohort, reflecting known site-specific variations in DCS protocols worldwide.^18,69^ This variation helped to assess the generalizability of our findings across differing DCS paradigms. At the original site (UCSF), but with a different surgeon, DCS was performed using bipolar electrodes using trains of 500-microsecond pulses delivered at 50 Hz. Each stimulation train lasted 2 seconds, and the current intensity ranged from 2–10 mA.^24,70,71^ At the secondary site (University of Michigan), direct cortical stimulation was delivered using a bipolar electrode with 5 mm spacing between contacts. Stimulation consisted of 1-millisecond biphasic square-wave pulses delivered in trains at 60 Hz, with a current intensity range of 2–4 mA across the series.^72^

### Intraoperative Subdural Electrocorticography

Following cortical exposure and prior to tumor resection, subdural electrocorticography (ECoG) recordings were obtained at a sampling rate of 4800 Hz. Depending on the extent of the craniotomy and the availability of materials, recordings were performed using either low-density (4×5 electrodes, 10 mm spacing) or high-density (8×12 electrodes, 5 mm spacing; or 10×10 electrodes, 3 mm spacing) subdural grids. For broader cortical coverage, up to two grids were used concurrently. After electrode placement, impedance testing was conducted to verify the integrity of the electrode contacts and ensure adequate contact with the cortical surface.

To minimize recording artifacts, all operating room personnel were instructed to refrain from unnecessary verbal communication and to turn off non-essential equipment (e.g., suction devices).^4^ Participants first completed a 3-minute eyes-closed resting-state baseline, during which they were asked to focus on their breathing. As described in our prior studies, this was followed by a visual confrontation naming (picture naming) task. Stimuli were presented on a 15-inch laptop (60 Hz refresh rate) placed approximately 30 cm from the participant. A custom MATLAB script built with PsychToolbox 3 (http://psychtoolbox.org/) displayed a single block of 48 unique, colored line drawings depicting common animals or objects. Participants were asked to vocalize the single word corresponding to each drawing. Trials were advanced manually by a clinician upon patient response, or automatically after 6 seconds if no response was given. During each picture naming trial, both the stimulus onset and speech onset times were recorded for downstream analysis.^3,4^

### Electrocardiography Preprocessing and Referencing

Picture-naming and resting-state ECoG recordings were initially imported into EEGLAB (MATLAB). Electrode channels were visually inspected, and channels exhibiting excessive noise were excluded. The remaining electrodes were re-referenced using a common average reference (CAR), wherein the mean signal across all retained electrodes was subtracted from each channel to reduce spatially widespread noise.^73^

### Grid Localization, Alignment of Electrodes to DCS mapped sides, and Creation of Patient-Specific Tumor Masks

Individual electrodes were aligned to DCS-mapped regions using intraoperative photographs, which were co-registered via an OpenCV-based script. To further refine localization, we utilized BrainTRACE (Brain Tumor Registration and Cortical Electrocorticography), a MATLAB tool developed by our group, to accurately project both intraoperative ECoG electrodes and DCS-mapped regions onto patient-specific pial surface reconstructions derived from preoperative MRIs.^74^ Alignment of DCS regions to electrode positions was additionally verified within BrainTRACE.

To create tumor masks, lesions were manually or semi-automatically segmented using 3D Slicer (https://www.slicer.org/) by a co-author blinded to the DCS regions. An abnormal FLAIR signal was used to delineate lesion boundaries across gliomas, maximizing the inclusion of radiographically identified infiltrative disease.

We, then, confirmed that all DCS-mapped regions were located within or adjacent to FLAIR hyperintense regions by exporting their stereotactic coordinates (x, y, z) from BrainTRACE and computing the Euclidean distance to the nearest tumor voxel from each patient’s tumor mask. For subsequent analyses, each DCS+ electrode was pair-matched to the nearest DCS– electrode within the same gyrus in each patient, enabling patient-specific, anatomically controlled comparisons. Examples of pair-matching DCS+ and nearest DCS– electrodes are illustrated in Extended Data Fig. 2c and d.

To visualize the locations of DCS+ and DCS– regions, we provide MNI maps (Fig. 1c, d and Fig. 2d, i) that coalesce regions across patients. Since these regions were initially localized on patient-specific pial surfaces using BrainTrace and subsequently transformed into MNI space, they should be interpreted as approximate, with possible minor deviations from their true anatomical locations.

Next, tumor masks were normalized for overlay and group-level visualization (Extended Data Fig. 2a-b). Using the Clinical Toolbox in SPM12 (https://www.fil.ion.ucl.ac.uk/spm/software/spm12/), each T2-weighted FLAIR-derived lesion mask was registered to the corresponding anatomical T1-weighted image. The T1-weighted images were then normalized to the standardized Montreal Neurological Institute template (MNI152). The resulting transformation matrix was applied to each participant’s lesion mask to bring it into MNI space. Normalized tumor volumes were then visualized in MNI152 space using MRIcroGL (www.nitrc.org).

### Task (Picture Naming) Event Related Spectral Perturbations (ERSPs)

Picture-naming recordings were down sampled to 600 Hz. Only correct trials were included. To calculate ERSPs, each trial was epoched separately for each electrode using MNE-Python and NeuroDSP.^75,76^ For all cortical regions—except the temporal lobe, due to its distinct response dynamics—epochs were centered around speech onset, spanning from −1 to +1 seconds. Baseline correction was performed using a variance-stabilizing transformation relative to the −1.2 to −1.0 s pre-speech window, followed by mean baseline subtraction. For temporal lobe electrodes, epochs were instead aligned to stimulus onset, with baseline correction from −0.2 to 0 seconds before stimulus onset. To compute broadband spectrograms (2–150 Hz), Morlet wavelets were used with a frequency-dependent number of cycles (3 to 50) increasing from low to high frequencies. For high gamma time series, wavelets were similarly applied but restricted to the 70–150 Hz range, with a higher number of cycles (30 to 50) to enhance frequency resolution.^77^

### Task-based comparison DCS+ and DCS− High Gamma Neuronal Time Series

The full patient cohort was divided into two groups based on tumor pathology: IDH-mutant (including oligodendrogliomas and astrocytomas) and IDH-wildtype (glioblastomas). Event-related spectral perturbations (ERSPs) in the high-gamma band (70–120 Hz)^27^ were computed by averaging spectral power at each time point from −1.0 to +1.0 seconds relative to speech onset across trials. To evaluate differences in high-gamma activation between DCS+ and DCS– electrodes, time series were averaged across trials and electrodes within each group and compared at each time point using Welch’s t-tests, with false discovery rate (FDR) correction for multiple comparisons. To determine whether each group’s high-gamma activity exceeded baseline levels, paired t-tests were performed between the baseline window (−1.2 to −1.0 s) and each post-stimulus time point, also FDR-corrected. These comparisons were conducted for all DCS+ and DCS– electrodes as well as for anatomically pair-matched DCS– electrodes.

For electrode-level summary metrics, average high-gamma power was computed from −0.5 to +0.5 seconds relative to speech onset. Group-level comparisons were then performed across all cortical regions and within the inferior frontal gyrus (IFG), with statistical significance assessed using two-tailed linear mixed-effects models controlling for patient-level random intercepts.

### Neuronal Encoding: High Gamma Entropy and Representational Similarity Analyses

High gamma power event-related spectral perturbation (ERSP) time series were computed for each trial and electrode as described above. For entropy analyses, ERSPs were grouped by DCS status (DCS+ vs. DCS–) within each patient cohort (IDH-mutant and IDH-wildtype). Shannon entropy was calculated across electrodes at each time point, either using all available DCS– electrodes or using pair-matched DCS– electrodes within the same gyrus as each DCS+ electrode.^4,34^ Comparisons between DCS+ and DCS– regions were performed using Welch’s t-tests.

For representational similarity analysis (RSA), only DCS+ electrodes and their pair-matched DCS– counterparts within the same gyrus were analyzed.^35^ Frontal and temporal electrodes were evaluated separately. Each electrode’s high gamma ERSP time series was tested for representational similarity with stimulus-level linguistic features. For frontal lobe electrodes, RSA was performed using both word frequency and semantic similarity, the latter derived from Global Vectors for Word Representation (GloVe) embeddings (300 or 50 dimensions).^78^ Temporal lobe electrodes were tested only with semantic similarity. Representational similarity was quantified using pairwise Pearson correlation between trials, and statistical significance was determined using nonparametric permutation testing (10,000 label shuffles per electrode). RSA was conducted independently for each patient, and results were aggregated across patients by DCS status.

### Decoding Word Frequency from High Gamma Activity

We utilized high-gamma ERSP time series to train multivariate linear regression models that predict the z-scored word frequency of spoken words, derived from the wordfreq Python package (version 3.1.1), based on a large language corpus. Models were trained and evaluated using leave-one-trial-out cross-validation within each patient. For each held-out trial, the model was trained on the remaining trials and used to predict word frequency from high gamma features. Prediction accuracy was quantified by trial-level root mean squared error (RMSE) between predicted and actual values. Decoding performance was grouped by electrode type (DCS+ vs. pair-matched DCS–). Analysis was restricted to frontal lobe electrodes. Statistical differences between groups were assessed using linear mixed-effects models, which included patient as a random effect to account for intra-patient variability. To characterize the significance of the decoders, we also performed 10 shuffles within each leave-one-out cross-validation. We compared the true decoding performance to the shuffled performance using linear mixed effects models that account for patient-level differences.

### Power spectral and aperiodic analysis of resting-state signals from DCS+ and DCS− regions

Resting-state data were downsampled to 1,200 Hz. These recordings were subsequently imported into Python and analyzed using MNE-Python.^75^ To suppress powerline noise and its harmonics, we applied a notch filter at 60 Hz, 120 Hz, 180 Hz, and 240 Hz using a zero-phase finite impulse response (FIR) filter, which attenuates narrowband artifacts while preserving broadband spectral content.^73^ To ensure consistent spectral representation across electrodes, power spectra were interpolated to a common frequency axis prior to spectral decomposition.^43^ Power spectral density (PSD) was estimated for each electrode using Welch’s method, with data segmented into 3-second epochs and 50% overlap. PSD was computed across canonical frequency bands: Delta (0.2–3.99 Hz), Theta (4–7.99 Hz), Alpha (8–12.99 Hz), Beta (13–29.99 Hz), Gamma (30–69.99 Hz), High Gamma (70–150 Hz), and a broadband Full Spectrum (0.2–150 Hz). Average power within each frequency band was then computed for each electrode.

To quantify broadband deviations, we first characterized the aperiodic (1/f-like) component of the power spectrum by applying the Fitting Oscillations and One-Over-F (FOOOF) algorithm (fooof 1.1.0 Python package).^44^ The model was initialized with parameters selected to capture physiologically relevant oscillatory structure: peak width limits of 0.5–12.0 Hz, a minimum peak height of 0.05 (in log power units), a peak threshold of 2.0 standard deviations above the aperiodic background, and a fixed aperiodic mode. FOOOF was applied to the interpolated spectra for each electrode, first over the full 2–150 Hz broadband range. Finally, the aperiodic fit was subtracted from the PSD in linear space, isolating periodic (oscillatory) contributions for subsequent analysis.

Power spectral density curves were constructed to examine broadband differences between DCS+ and DCS– electrodes.^43^ To statistically compare spectral features, linear mixed-effects models were fit with patient as a random effect. These models assessed narrowband power across frequency bands from Theta to High Gamma, broadband deviations from the aperiodic slope, and the slope exponent of the aperiodic (1/f) component within the high gamma range.

### Classification of DCS Status Using Resting State Spectral Features

To assess whether power spectral features could distinguish language-positive (DCS+) from language-negative (DCS-) electrodes, we trained machine learning classifiers (least absolute shrinkage and selection operator (LASSO) regression, ridge regression, elastic net regression, and logistic regression) on PSD profiles. PSD estimates were computed as described above and used as input features. For internal validation, data from patients with both DCS+ and DCS– electrodes were randomly split into training (80%) and testing (20%) sets. Classifier performance was assessed on held-out electrodes within this internal cohort. To assess statistical significance, we performed non-parametric permutation testing by randomly shuffling the training labels 1,000 times and recalculating the classifier accuracy on the unshuffled test set to generate a null distribution.

For external validation, models were trained on all internal patients with both DCS+ and DCS– regions and tested on an independent external cohort of patients not included in the training set. For each held-out electrode in the external set, we evaluated both the binary prediction and the associated SoftMax probability score to assess classifier confidence, recognizing that real-world outcomes are rarely strictly binary.

