## Extended Data Figures 1-5 for "Electrocorticographic Detection of Speech Networks in Glioma-infiltrated Cortex"

Extended data Figure 1

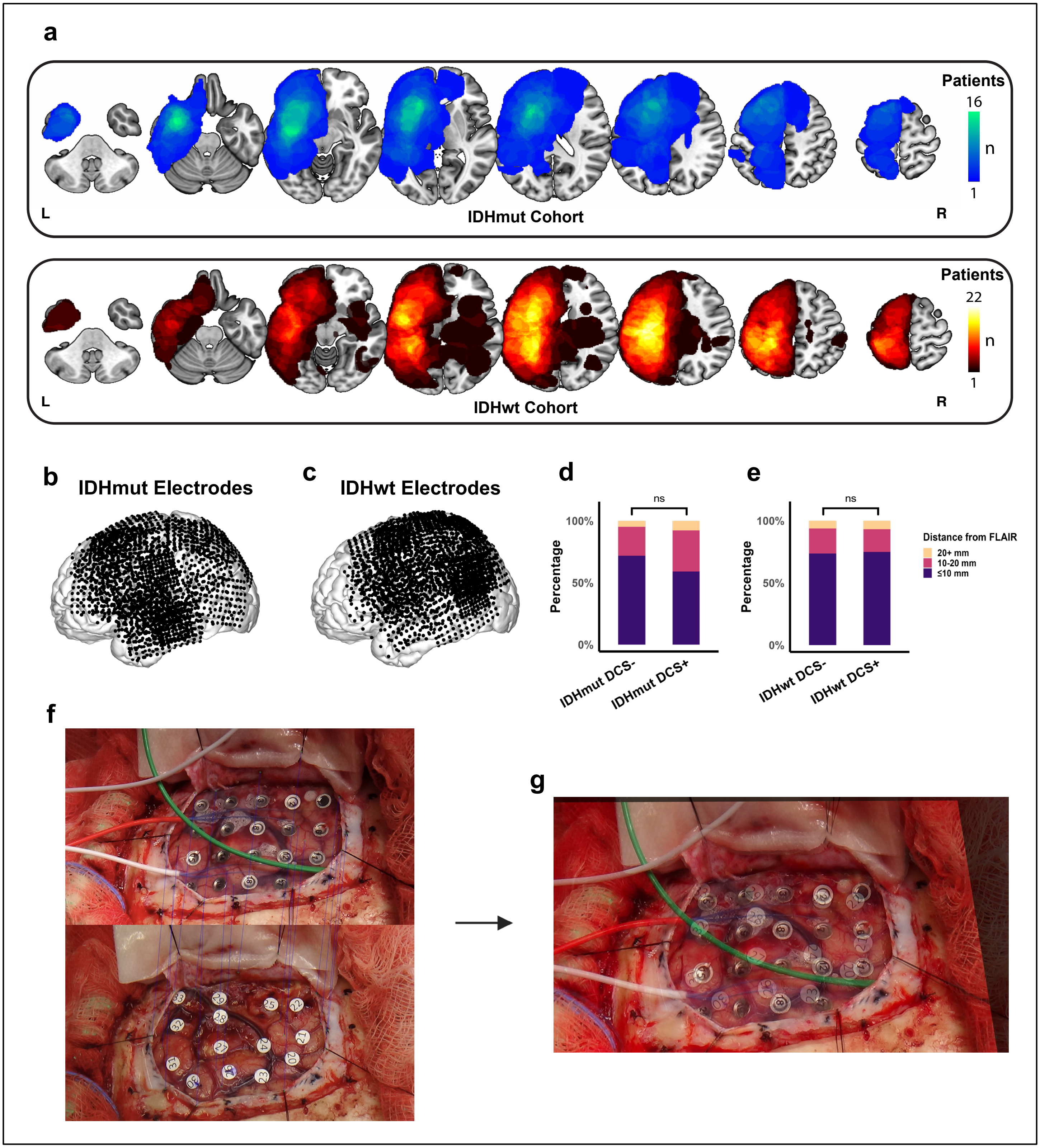

**a,** Representative axial MNI brain slices showing tumor masks across IDH-mutant cohort (36 patients, top blue) and IDH-wildtype cohort (45 patients, bottom red). **b-c**, Electrode grids, with each dot representing a single electrode, projected into MNI space for all patients in the **b**, IDH-mutant, and **c**, IDH-wildtype cohorts. **d-e,** DCS+ and DCS– sites do not differ significantly in distance from the FLAIR hyperintensity edge in either the **d**, IDH-mutant (p=0.23) or **e**, IDH-wildtype cohorts (p=0.95). Statistical significance was assessed using a Chi-squared test across distance bins. **f**, Example of an open computer vision (CV)-based script output that allows users to identify regions of interest by aligning an image of the electrode grid (top) with the corresponding DCS mapping (bottom), yielding **g**, a co-registered image that facilitates accurate assignment of electrodes to DCS sites.

**Extended data Figure 2**

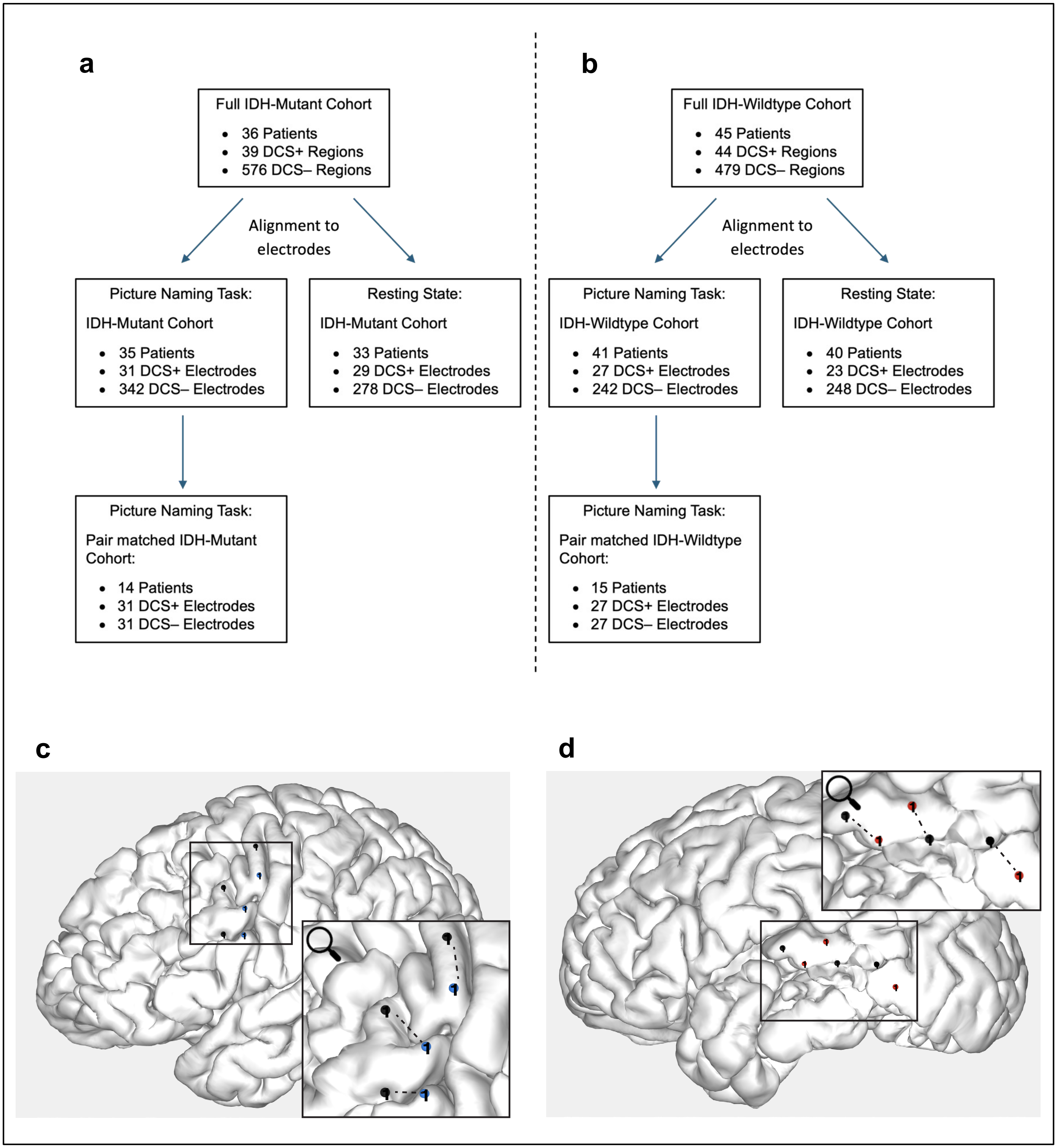


**a-b**, Flow chart demonstrating how total mapped sites were aligned to electrodes and included in subsequent picture naming and resting state analyses for **a**, IDH-mutant and **b**, IDH-wildtype cohorts. The number of patients and electrodes may vary by task due to differences in electrode processing and patient participation. **c–d**, Representative examples of pair-matching on patient-specific pial maps. **c**, IDH-mutant patient with DCS+ sites shown in blue. **d**, IDH-wildtype patient with DCS+ sites shown in red. In both panels, pair-matched DCS– sites are shown in black.

**Extended data Figure 3**

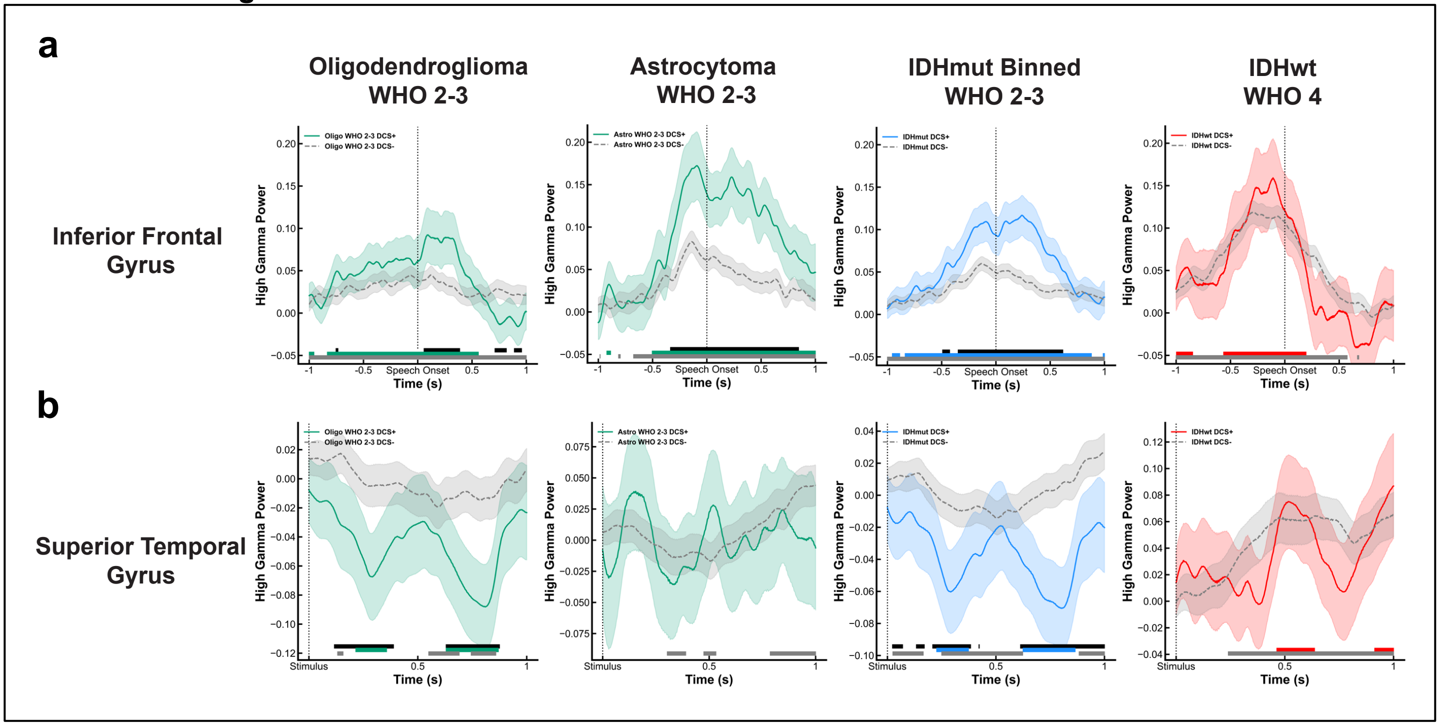


**a**, High-gamma power time series, aligned to speech onset, from inferior frontal gyrus (IFG) DCS+ and DCS– sites demonstrate significant differences (black bar) in IDH-mutant glioma cohorts but not in IDH-wildtype glioblastoma. From left to right, tracings are shown for Oligodendroglioma (18 patients; 8 DCS+ and 33 DCS– electrodes), Astrocytoma (17 patients; 6 DCS+ and 29 DCS– electrodes), combined IDH-mutant (35 patients; 14 DCS+ and 62 DCS– electrodes), and IDH-wildtype glioblastoma (40 patients, 8 DCS+ and 46 DCS– electrodes). **b**, High-gamma power time series, aligned to prompt onset, from superior temporal gyrus (STG), DCS+ and DCS– sites demonstrate significant differences (black bar) in IDH-mutant glioma cohorts (except Astrocytoma alone due to only 1 DCS+ site in STG) but not in IDH-wildtype glioblastoma. From left to right, tracings are shown for Oligodendroglioma (7 patients; 5 DCS+ and 18 DCS– electrodes), Astrocytoma (8 patients; 1 DCS+ and 22 DCS– electrodes), combined IDH-mutant (15 patients; 6 DCS+ and 40 DCS– electrodes), and IDH-wildtype glioblastoma (11 patients, 7 DCS+ and 25 DCS– electrodes). In panels **a**, **b**, solid lines represent the cohort-averaged time series across trials and electrodes; shaded areas denote standard error of the mean (SEM). Time series were grouped by DCS status and compared at each time point using Welch's *t*-tests with false discovery rate (FDR) correction for multiple comparisons. Time points with significant differences are marked by black bars above the x-axis. Colored bars denote time points where each cohort's High-gamma power was significantly greater than its own baseline (paired *t*-test, FDR corrected).

**Extended data Figure 4**

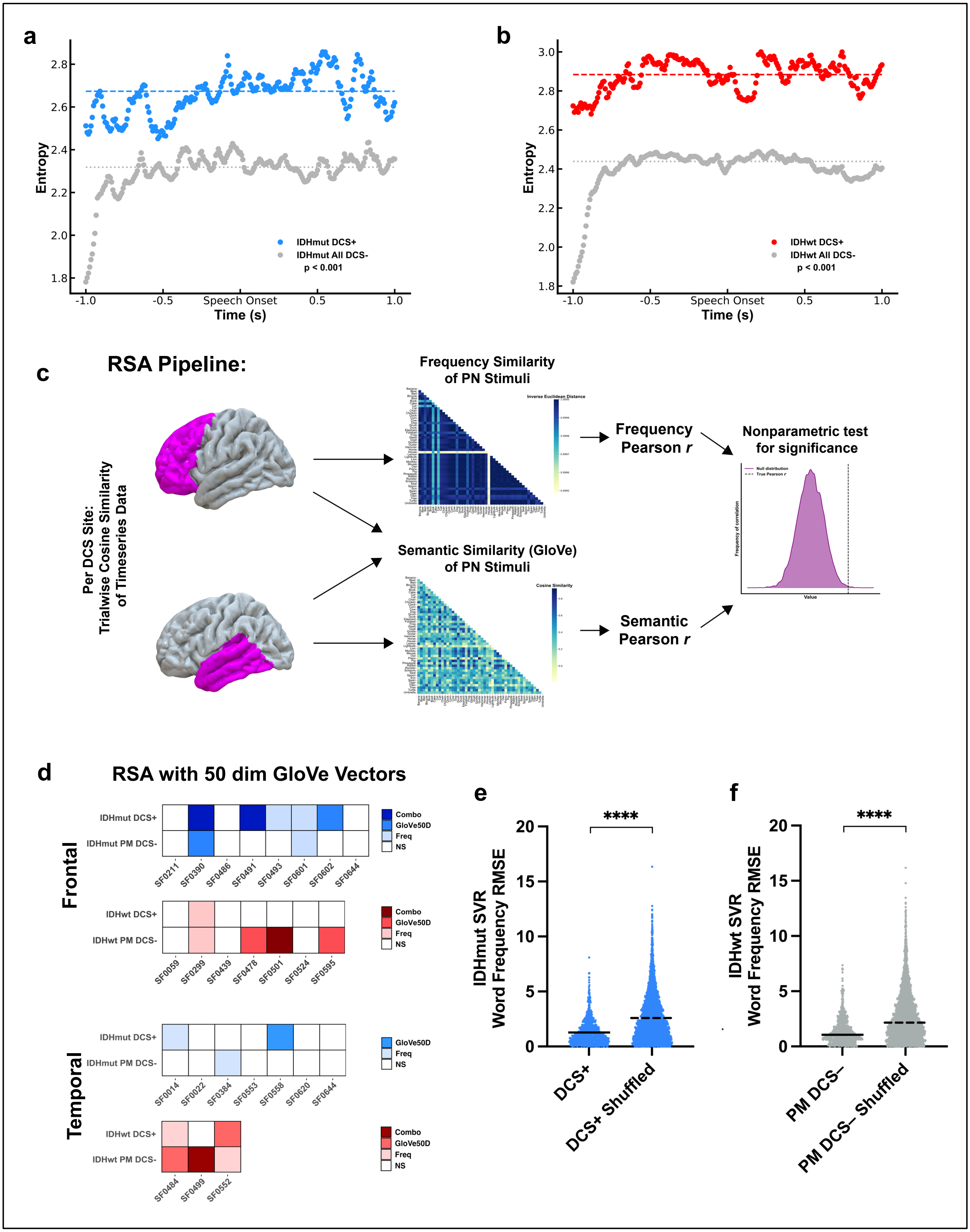

**a-b**, High gamma power ERSP entropy in **a,** IDH-mutant (a) and **b,** IDH-wildtype (b) cohorts, using all DCS– electrodes rather than only pair-matched controls, as shown in **Fig. 3 a-b**. DCS+ sites continued to exhibit significantly higher entropy across time in both IDH-mutant ($p < 0.001$) and IDH-wildtype ($p < 0.001$) cohorts. **c,** Schematic of the representational similarity analysis (RSA) pipeline. Frontal and temporal lobe electrodes were tested for representational similarity with both word frequency and semantic (GloVe) embeddings. Statistical significance was assessed via 10,000-label shuffling for nonparametric testing. **d,** Replication of analysis from **Fig. 3c** using 50-dimensional GloVe embeddings. Frontal and temporal lobe electrodes were tested for both word frequency and semantic similarity. In the IDH-mutant cohort, DCS+ electrodes were more likely to exhibit significant RSA within the same patient. This pattern was not observed in the IDH-wildtype cohort. **e**, In the IDH-mutant cohort, decoding with DCS+ electrodes from **Fig. 3f** was significantly better than chance, as assessed by comparison to their shuffled controls (DCS+ $1.27 \pm 1.14$ vs DCS+ shuffle: $2.69 \pm 2.38$, $p < 0.001$). f, In the IDH-wildtype cohort (Supp. Fig. 3f), decoding with DCS– electrodes from **Fig. 3e** was also significantly better than chance (DCS– $1.48 \pm 1.37$ vs DCS– shuffle: $2.48 \pm 2.25$, $p < 0.001$). **e-f**, Statistical significance was assessed using linear mixed-effects models accounting for patient level differences. Horizontal black lines represent cohort means.

**Extended data Figure 5**

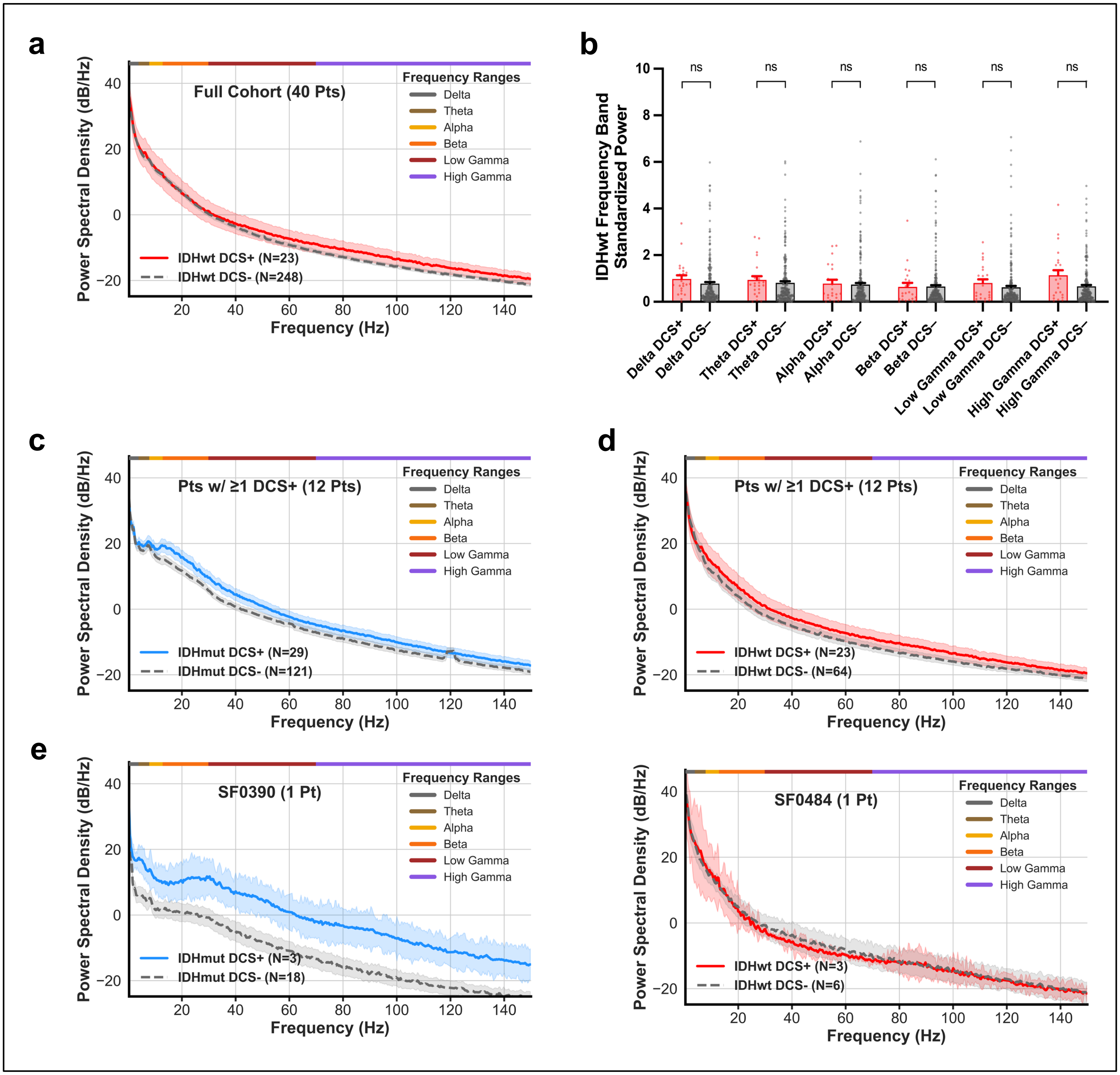


**a,** Power spectral density (PSD) curves (left) from resting state recordings demonstrate clear separation of IDH-wildtype cohort (40 patients), DCS+ language sites (N=23), and DCS– language sites (N=248). Shaded regions demonstrate the subgroup's standard error of the mean (SEM). **b**, Linear mixed-effects model-based analysis of individual frequency ranges (Delta to High Gamma) demonstrates no significant differences between IDH-wildtype DCS+ and DCS– sites, while controlling for patient-level differences (all p>0.05). **c**, PSD curves of IDH-mutant DCS+ and DCS– sites maintain the separation observed in **Fig. 3a** when only including the subset of patients with at least one DCS+ site. **d**, PSD curves of IDH-wildtype DCS+ and DCS– sites, when only including the subset of patients with at least one DCS+ site, do not separate, consistent with **Supp. Fig. 4a**. **e**, Representative PSDs of a single IDH-mutant patient (left) and a single IDH-wildtype patient (right).
